# Multi-modal single cell analysis reveals brain immune landscape plasticity during aging and gut microbiota dysbiosis

**DOI:** 10.1101/2020.07.15.205377

**Authors:** Samantha M. Golomb, Ian H. Guldner, Anqi Zhao, Qingfei Wang, Bhavana Palakurthi, Jacqueline Lopez, Kai Yang, Siyuan Zhang

## Abstract

The brain contains a diverse array of immune cell types. The phenotypic and functional plasticity of brain immune cells collectively contribute to brain tissue homeostasis and disease progression. Immune cell plasticity is profoundly influenced by local tissue microenvironment cues and systemic factors. Yet, the transcriptional mechanism by which systemic stimuli, such as aging and gut microbiota dysbiosis, reshape brain immune cell plasticity and homeostasis has not been fully delineated. Using Cellular Indexing of Transcriptomes and Epitopes by sequencing (CITE-seq), we analyzed compositional and transcriptional changes of the brain immune landscape in response to aging and gut dysbiosis. We first examined the discordance between canonical surface marker-defined immune cell types (Cell-ID) and their transcriptome signatures, which suggested transcriptional plasticity among immune cells despite sharing the same cell surface markers. Specifically, inflammatory and patrolling Ly6C+ monocytes were shifted predominantly to a pro-inflammatory transcriptional program in the aged brain, while brain ILCs shifted toward an ILC2 transcriptional profile. Finally, aging led to an increase of ILC-like cells expressing a T memory stemness (T_scm_) signature in the brain. Antibiotics (ABX)-induced gut dysbiosis reduced the frequency of ILCs exhibiting T_scm_-like properties in the aged mice, but not in the young mice. Enabled by high-resolution single-cell molecular phenotyping, our study revealed that systemic changes due to aging and gut dysbiosis prime the brain environment for an increased propensity for neuroinflammation, which provided insights into gut dysbiosis in age-related neurological diseases.

**Manuscript Summary:** Golomb *et al.* performed Cellular Indexing of Transcriptomes and Epitopes by sequencing (CITE-seq) on immune cells from the brains of young and aged mice with and without antibiotics-induced gut dysbiosis. High resolution, single cell immunophenotyping enabled the dissection of extensive transcriptional plasticity of canonically identified monocytes and innate lymphoid cells (ILCs) in the aged brain. Through differential gene expression and trajectory inference analyses, the authors revealed tissue microenvironment-dependent cellular responses influenced by aging and gut dysbiosis that may potentiate neuroinflammatory diseases.

**Graphical Abstract:** 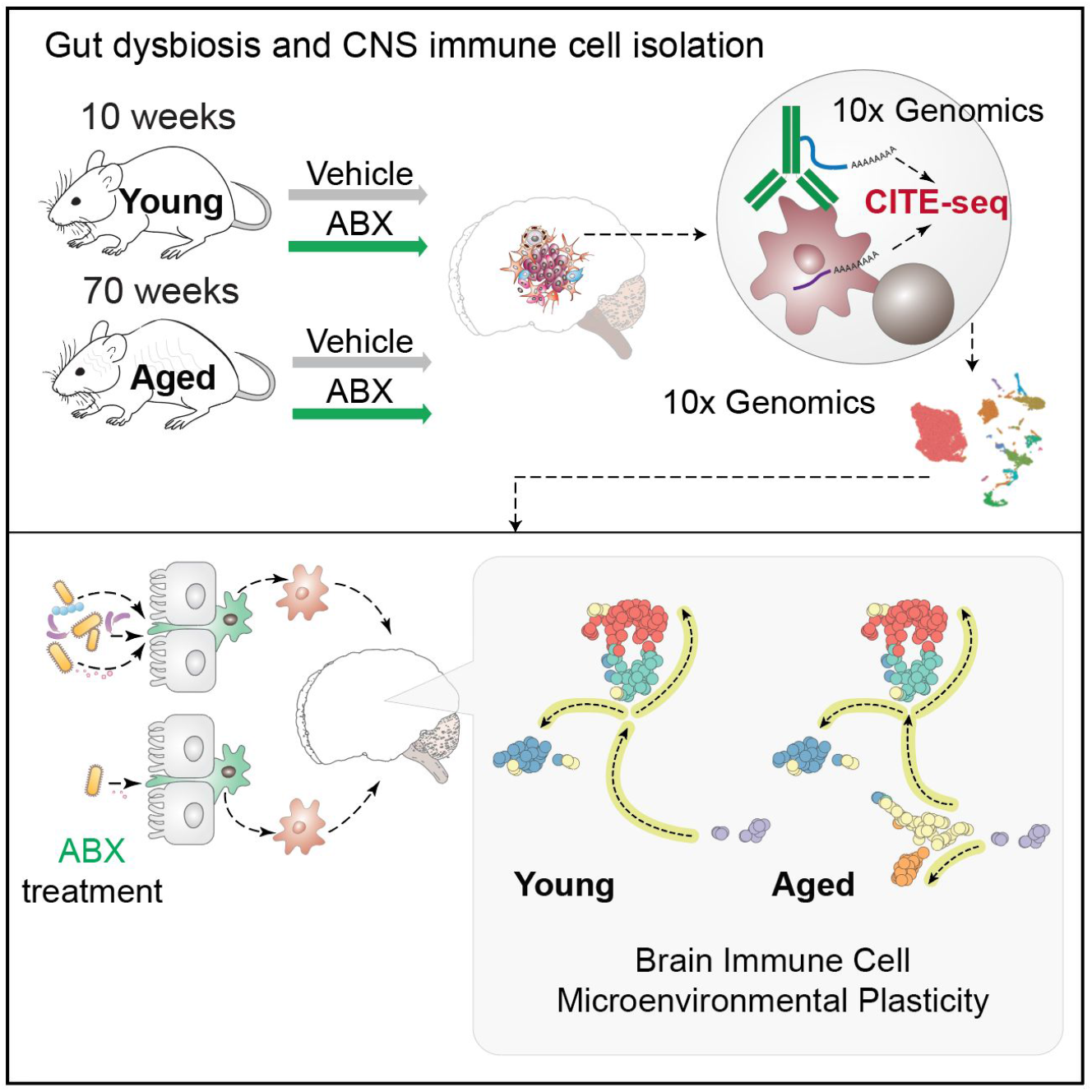

## INTRODUCTION

The brain has been long considered an immune-privileged site. However, it has become increasingly clear that the level of immune-privilege varies within the steady-state brain depending on age and neurological health (Mrdjen et al., 2018). Recent studies have revealed extensive compositional and transcriptional heterogeneity of the brain resident and peripherally derived immune cells in the central nerve system (CNS). Such heterogeneity collectively manifests a highly dynamic and plastic CNS immune milieu. CNS resident and peripheral immune cells of distinct hematopoietic lineages act in concert to maintain CNS homeostasis. Dysregulation of such homeostasis significantly contributes to age-related neurodegenerative diseases, neuroinflammation, and brain tumors (Consortium et al., 2019; Dulken et al., 2019; Keren-Shaul et al., 2017; Mrdjen et al., 2018; Quail and Joyce, 2017; Ximerakis et al., 2019).

Aging is an influential systemic factor contributing to the decline of tissue function and the functionality of the immune system. The aging process leads to natural perturbations of microbial composition, namely gut microbiota dysbiosis, which is believed to contribute to systemic inflammation (Franceschi et al., 2018; Langille et al., 2014; Levy et al., 2017; O’Toole and Jeffery, 2015; Thaiss et al., 2016). The gut microbiota engages in crosstalk with both innate and adaptive immune systems through either direct engagement of the mucosal innate immune system or commensal derived metabolites (Belkaid and Hand, 2014). Gut microbiota dysbiosis significantly alters the circulating metabolites and plasma cytokine composition, leading to dysregulation of the peripheral immune system (Arpaia et al., 2013; Bachem et al., 2019; Lehallier et al., 2019). Intriguingly, beyond peripheral immunity, studies have suggested that gut microbiota dysbiosis also indirectly regulates CNS immunity and neuroinflammation through microbiota-derived signaling molecules (Dinan and Cryan, 2017; Erny et al., 2015; Ma et al., 2019; Sampson et al., 2016).

Maintaining homeostasis of CNS immunity under systemic change requires CNS resident and CNS infiltrating immune cells to manifest substantial functional plasticity in response to both microenvironmental and systemic changes, such as aging and dysbiosis. In particular, there are potentially two forms of plasticity among immune cells: 1) intra-lineage cell plasticity, referring to the phenotypic changes within a given cell lineage and 2) trans-lineage cell plasticity, referring to the potential transdifferentiation between canonically defined lineages (Gagliani et al., 2015; Galli et al., 2011; Meister and Ferrandon, 2011). At both the compositional and single-cell transcriptome levels, age-associated compositional shifting of various CNS resident immune cells, such as microglia and border associated macrophages (BAMs), and peripheral immune cells have been observed (Dulken et al., 2019; Mrdjen et al., 2018; Ximerakis et al., 2019). However, the age-related immune plasticity of peripherally-derived brain infiltrating innate immune cells, such as Ly6C+ monocytes, as well as innate lymphoid cells (ILCs), have not been fully delineated.

The recent advances in multi-modal single-cell analysis platforms, namely Cellular Indexing of Transcriptomes and Epitopes by sequencing (CITE-seq), has enabled a comprehensive single-cell immunophenotyping by connecting canonical immune cell lineage identity to the cellular status through transcriptome readout (Stoeckius et al., 2017). In this study, we sought to map brain immune cell plasticity in response to systemic perturbations of aging and gut dysbiosis. Using CITE-seq, our study provides a single cell level characterization of the compositional and transcriptional plasticity of brain immunity, as exemplified in the transcriptome plasticity among inflammatory/patrolling Ly6C+ monocytes and CNS-associated ILCs. Revealing such immune cell plasticity during aging and gut dysbiosis sheds light on critical components governing brain immunity in aging and the onset of age-related neurodegenerative disease.

## RESULTS

### CITE-seq delineates the global immune cell diversity in the brain

The brain contains a diverse immune cell population as identified through CyTOF or scRNA-seq analysis (Mrdjen et al., 2018; Ximerakis et al., 2019). Here, we employed CITE-seq analysis to simultaneously profile the surface marker expression and transcriptional status of individual brain immune cells. To further explore dynamics of the brain immune cell landscape in response to systemic changes imposed by aging and gut microbiota dysbiosis, we isolated whole brains from twelve young adult (13 weeks old, human equivalent ~20 years old) and twelve aged (73 weeks old, human equivalent ~56 years old) mice for CITE-seq analysis (**Fig. 1A**). Antibiotics treated (ABX) and control groups received an antibiotic cocktail or vehicle by oral gavage daily for three weeks to induce gut microbiota depletion. At the completion of the three week treatment, fecal samples were processed for 16S analysis, and brains were collected, digested, and enriched for immune cells. The anti-bacterial efficacy of the antibiotic treatment strategy as well as naturally occurring gut dysbiosis in aged animals, were confirmed by Illumina sequencing of the 16S rRNA V3-V4 regions of microbial DNA extracted from mouse fecal pellets. 16S sequencing analysis revealed that ABX treatment caused significant shifting in bacterial phyla composition and reduction in alpha diversity measured by Shannon’s diversity index (**Supplementary Fig. 1A-B**). With ABX treatment, young and aged mice shifted from *Bacteroidetes* and *Firmicutes* being the predominant bacterial phyla, respectively, to an enrichment of *Proteobacteria* in both ages (**Supplementary Fig. 1A**). The brain immune cells were stained with a panel of 31 antibodies (**Supplementary Table 1**) before 10X Genomics Chromium Single Cell Gene Expression analysis.

**Figure 1:**
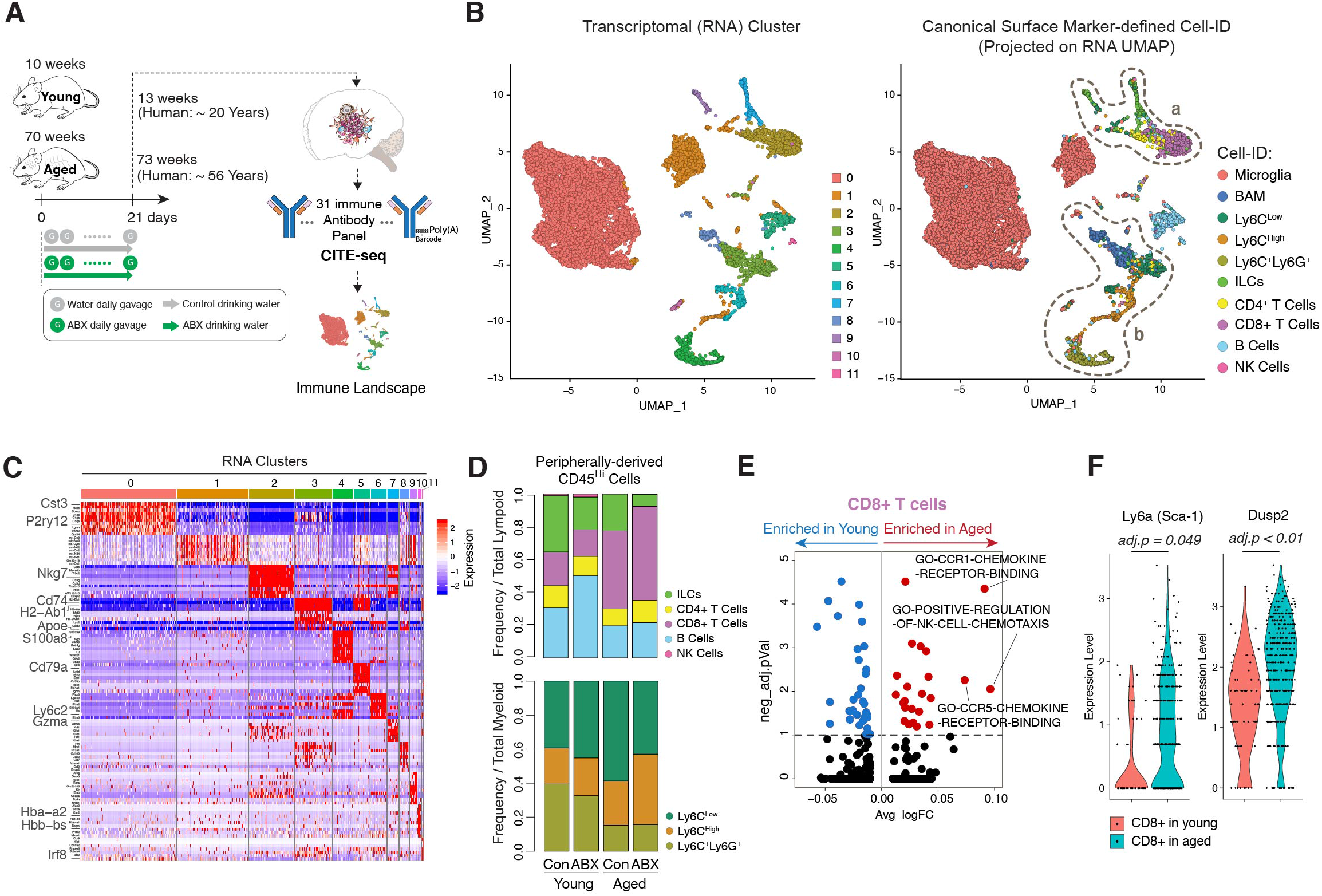
CITE-seq delineates the global immune cell diversity in the brain. A) Schematic of the experimental approach. B) (left) Single cells projected onto UMAP. Cells are color-coded by transcriptional cluster ID. (right) UMAP clustering as in the left plot but cells are color-coded by canonical cell ID. Dashed circle (a) encompasses peripheral lymphoid cell populations and dashed circle (b) encompasses non-microglia myeloid cells. C) Heatmap of genes organized by transcriptional clusters. D) Stacked bar charts of the proportions of peripheral lymphoid cells (top) and peripheral myeloid cells (bottom) for each experimental condition. E) Volcano plot showing differentially enriched gene pathways in CD8+ T cells from aged mice compared to young mice. F) Violin plots of *Ly6a* and *Dusp2* expression level in CD8+ T cells from young and aged mice.

To analyze the transcriptional similarities and profiles among the immune cells, single cells were clustered using a principal component analysis (PCA)-based approach and projected by Uniform Manifold Approximation and Projection (UMAP) onto a two-dimensional plot (**Fig. 1B left**). Clustering analysis revealed 11 transcriptionally distinct clusters of cells present in all samples. Gene signatures for each cell cluster were generated by differential gene expression analyses comparing one cell type against the other cell populations present in the dataset (**Fig. 1C** and **Supplementary Table 2**). CITE-seq enabled us to perform a joint-analysis whereby we performed traditional canonical surface marker-based gating (Cell-ID) using CITE-seq antibodies (**Supplementary Fig. 2)**, and then simultaneously projected Cell-ID distribution on the RNA-based UMAP plot (**Fig. 1B, right**). Canonically identified cells segregated into different clusters that were generally mutually exclusive. However, it was noted that there was increased heterogeneity among the peripherally-derived lymphoid and myeloid populations (denoted by the dashed lines; **a** and **b**) (**Fig. 1B right** and **Supplementary Fig. 3A**). The majority (72%) of brain immune cells consisted of CNS native microglia (CD45^Low^,CD11b^+^,CD38^Low’^MHCII^Low^, *Tmem119^+^,Mrc1^-^*). Border associated macrophages (BAMs) were distinguished from microglia based on their relatively high expression of CD38, MHCII, and *Mrc1*, and lack of *Tmem119* expression and made up 3.9% of all collected immune cells (**Supplementary Table 3**) (Mrdjen et al., 2018).

Peripherally-derived CD45^Hi^ innate and adaptive immune cells were expectedly present at lower frequencies (24%) among all of the collected leukocytes, but exhibited variability in lymphoid and myeloid subset proportions among the four conditions (**Fig. 1D** and **Supplementary Table 3**). The CD45^Hi^ peripherally-derived innate immune cells were segregated into lymphoid and myeloid categories. Within the lymphoid compartment, there was more variable shifting among B cells (between young control and young ABX), CD8+ T cells (between young and aged), and ILCs (across all groups). Compared to young controls, young ABX mice experienced a slight increase in the proportion of B cells within the lymphoid compartment, from 2.6% in young control to 4.3% in young ABX. B cells were relatively stable in maintaining the RNA cluster 5 gene signature (*Cd79a, Ly6d* and *Iglc2*) but did display ABX-associated dispersion into other transcriptional clusters (RNA clusters 1, 3, 4 and 11). Dispersion away from the main cluster is represented by the slight decrease in canonically identified B cell abundance within RNA cluster 5 (young control: 75%, young ABX: 68%, aged control: 68%, aged ABX: 54%) (**Supplementary Table 3,** Cell-ID vs RNACluster Frequency).

T cells were identified by CD3 surface protein expression and *Thy1+/Itga2+/Klrb1-* at the RNA level (**Supplementary Figure 2,** gating strategy). The total T cell population was further segregated into CD4+ T cells (1.7% of leukocytes) and CD8+ T cells (5.7% of leukocytes) (**Supplementary Table 3**). Consistent with previous reports (Dulken et al., 2019), we noted an increased frequency of CD8+ T cells in aged control (8.3%) and aged ABX (11%) compared to young control (1.6%) and young ABX (1.6%) mice (**Fig. 1D** and **Supplementary Table 3**). We further explored the age-dependent transcriptional differences in the CD8+ T cell population using GSVA analysis. CD8+ T cells in aged mice were enriched in gene pathway components associated with immune cell chemotaxis including CCR1 chemokine receptor binding and positive regulation of NK cell chemotaxis (**Fig. 1E** and **Supplementary Table 4,** GSVA analysis). Of particular interest, CD8+ T cells from aged mice overexpressed *Ly6a* (avg log FC = 0.646, adj. p-value < 0.05), a stem cell marker, and *Dusp2* (avg log FC = 0.639, adj. p-value = 5.89E-05) (**Fig. 1F** and **Supplementary Table 2**). The DUSP family of phosphatases are described to be involved in regulating immune effector responses upon recognition of antigen. In particular for lymphocytes, upregulation of *Dusp2* is associated with engaging efficient clonal expansion (Lang and Raffi, 2019). The upregulation of numerous ribosomal proteins (*Rpl38, Rpl37, Rpl39*) as well as *Dusp2* (**Supplementary Figure 3B** and **Supplementary Table 2**), suggests that these T cells are actively proliferating or expanding within the aged brain (Dulken et al., 2019; Zhou et al., 2015). Young and aged mice with ABX-induced gut dysbiosis did not experience further changes in the expression of these genes (**Supplementary Fig. 3C** and **Supplementary Table 2**), suggesting the observed phenotype is primarily age-dependent.

CD45^Hi^ peripherally-derived innate myeloid cells were segregated from lymphoid cells on the basis of *Itga2* and *Thy1* expression (**Supplementary Fig. 2**). Ly6C+ myeloid cells, which were largely present in RNA clusters 1, 3, 4 and 6, globally expressed *Ifitm3, Lgals3* and *Vim* (**Fig. 1B-C** and **Supplementary Table 2**). Ly6C+ cells were further segregated into three subpopulations typically described as: 1) Ly6C^Hi^ monocytes (3.1% of leukocytes, expressing *Plac8, Fn1* and *S100a4*); 2) Ly6C^Lo^, patrolling monocytes (4.9% of leukocytes, expressing *H2-Aa, H2-Eb1* and *Cd74*), and 3) Ly6C^+^Ly6G^+^, neutrophils (2.5% of leukocytes, expressing *S100a9, S100a8, Retnlg* and *Lcn2*) (**Fig. 1C, Supplementary Tables 2** and **3**). Among these peripherally-derived myeloid cells, there was an age-associated shifting of Ly6C+Ly6G+ cells (neutrophils). Canonically identified neutrophils did not exhibit obvious signs of plasticity in the aging or gut dysbiosis contexts by maintaining the RNA cluster 4 gene signature (*S100a9, S100a8* and *Retnlg*) (young control: 99%, young ABX: 98%, aged control: 98%, aged ABX: 98%) (**Fig. 1C** and **Supplementary Table 3**). In comparison to aged control and aged ABX mice, there was an increased frequency of Ly6C^High^ and reduced abundance of Ly6C^Low^ cells in the aged ABX group (**Fig. 1D** and **Supplementary Table 3**).

### Aged brain is enriched for inflammation-prone brain resident myeloid cells

Among the CNS resident myeloid cells, we identified two major CD45^Lo^CD11b^+^ CNS native cell types - microglia (72% of leukocytes, expressing *Cx3cr1, Tmem119, P2ry12, Hexb*, and *Cst3*) and BAMs (3.9% of leukocytes, expressing *Cd74, Apoe, H2-Aa, H2-Ab1*, and *Mrc1*) (**Supplementary Table 3**). The canonically identified microglia largely segregated into two transcriptionally distinct clusters (RNA clusters 0 and 1) (**Fig. 2A-B**). Cell-ID Microglia predominantly expressed the RNA cluster 0 gene signature with an increased abundance of microglia within the RNA cluster 1 signature present in the aged control (15%) and even higher in aged ABX (17%) in contrast to young control (11%) and young ABX (10%) (**Fig. 2B top** and **Supplementary Table 3**). GSVA analysis revealed that cluster 1 is enriched with gene pathways involved in MHC-II protein binding, dendritic cell interaction, and macrophage cytokine production (**Fig. 2B bottom** and **Supplementary Table 4,** C1vsC0). Enrichment in these pathways suggests that microglia with the cluster 1 signature are in a more proliferative and pro-inflammatory state, which was further confirmed with the increase of mitochondrial genes (e.g. *mt-Co3*) and decreased expression of microglia homeostatic genes (e.g. *Trem2, Cst3*, and *Hexb*) (**Fig. 2C**). The slight increase of cluster 1 microglia population in both the aged brain and ABX treated groups implies gut dysbiosis increased the propensity for neuroinflammatory development in the aged brain.

**Figure 2:**
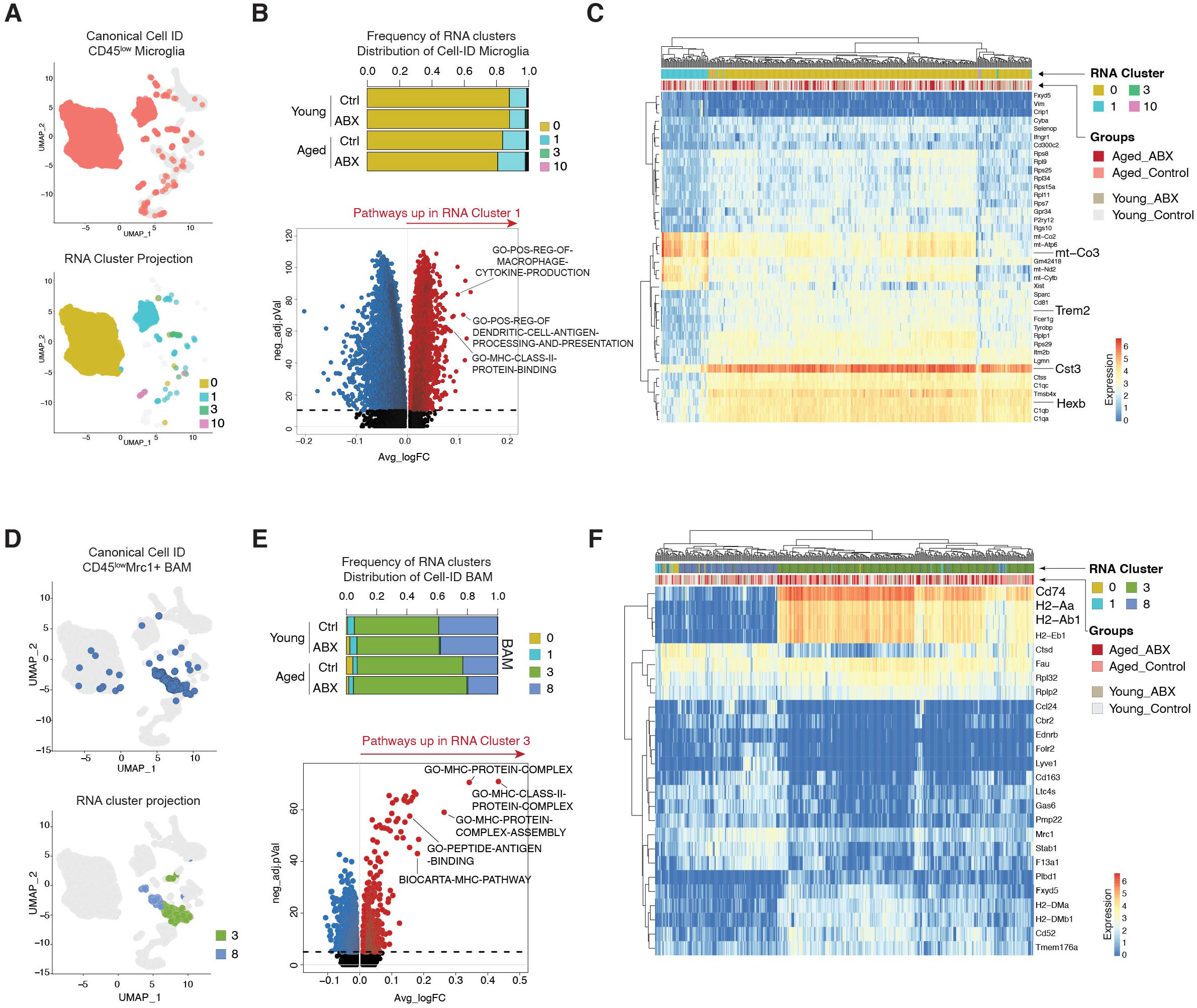
Aged brain is enriched for inflammation-prone brain resident myeloid cells. A) (top) UMAP clustering as in (1B) with Cell-ID microglia colored in pink (all other cells in gray). (bottom) UMAP clustering as in (1B) with Cell-ID microglia colored by transcriptional clusters. B) (top) Stacked bar charts of Cell-ID microglia within transcriptional clusters within each sample group. (bottom) Volcano plot showing differentially enriched gene pathways in RNA cluster 1 compared to RNA cluster 0. C) Heatmap with top 40 marker genes expressed in Cell-ID microglia. D) (top) UMAP clustering as in (1B) with Cell-ID BAMs only in blue (all other cells in gray). (bottom) UMAP clustering as in (1B) with Cell-ID BAMs colored by transcriptional cluster ID. E) (top) Stacked bar charts of Cell-ID BAMs within transcriptional clusters within each sample group. (bottom) Volcano plot showing differentially enriched gene pathways in RNA cluster 3 compared to RNA cluster 8. F) Heatmap with top 26 marker genes expressed in Cell-ID BAMs.

Cell-ID BAMs were predominantly present in RNA clusters 3 and 8 with only a small portion of cells expressing cluster 0/1 signature (**Fig. 2D-E**). BAMs with the RNA cluster 3 gene profile from the aged control and aged ABX were in higher proportion (70% and 75% respectively) compared to BAMs with RNA cluster 8 gene profile (23% and 20% respectively). On the other hand, BAMs in young control and young ABX groups were split ~ 60% to 40% between RNA cluster 3 and 8 gene profiles, respectively (**Fig. 2E top**). GSVA analysis revealed that RNA cluster 3 was enriched for gene pathways involved in MHC-II protein complex assembly and antigen peptide binding (**Fig. 2E bottom** and **Supplementary Table 4**). This gene signature resembles a transcriptional profile described for disease-associated BAMs in previous literature (Jordão et al., 2019). Marker genes for cluster 3 (*Cd74, H2-Aa*, and *H2-Ab1*) further support MHC-II pathway enrichment and potential antigen presentation activity of BAMs (**Fig. 2F** and **Supplementary Table 2**). Similar to microglia, the increased proportion of BAMs with the “disease-associated” signature in the aged mice suggests that BAMs in the aged CNS are shifting their transcriptome status to adapt to the inflammation-prone CNS tissue environment during aging.

### Innate Ly6C^High^ monocytes show microenvironmental-dependent plasticity in aged brain

Monocytes with high Ly6C surface protein expression (Ly6C^Hi^) are categorized as innate pro-inflammatory responders and exhibit plastic qualities in response to stimuli. Upon recruitment to sites of injury or infection, these cell types often mature into inflammatory macrophages while also secreting pro-inflammatory cytokines (Yang et al., 2014). Recruitment of these cell types and their maturation into pro-inflammatory responders can also cause tissue degradation and T cell activation (Yang et al., 2014). Compared to CNS-native myeloid cells, we observed relatively increased transcriptional plasticity among the Ly6C+ compartment in response to aging (**Fig. 3A**). In young mice, Cell-ID Ly6C^Hi^ cells largely centered around RNA cluster 6 core signature (**Fig. 3A left**). Interestingly, in aged control and aged ABX mice, Cell-ID Ly6C^Hi^ monocytes exhibited a more diverse transcriptome status, spreading across several RNA clusters (**Fig. 3A right**). Such transcriptional heterogeneity suggests a plastic nature of this cell type in response to the aging process and microbiota depletion. To gain additional insights into the plasticity of these cells, we subsetted canonically identified Ly6C^Hi^ cells and re-clustered the cells into four transcriptionally distinct subclusters with a varying frequency between young and aged (**Fig. 3B, top**). Subcluster 0 expressed inflammatory genes, including *Ifitm3, Lyz2*, and *Cxcl2*. Subcluster 3 expressed proliferation markers, *Mki67* and *Top2a* (**Fig. 3B bottom** and **Supplementary Table 2**). In the aged mice group, there was an increased frequency of Ly6C^Hi^ subcluster 1 and emergence of subcluster 2 with high expression of *Cstg, Mpo, Elane* (**Fig. 3B bottom**). RNA velocity analysis of the reclustered Ly6C^Hi^ cells based on RNA splicing showed a prevalent continuous pattern of cell velocity vector field (arrows) connecting subclusters, suggesting the subclusters are closely related with potentially continuous cellular statuses (**Supplementary Figure 4A**). We next employed a trajectory inference analysis, Slingshot, through the Dyno package (https://dynverse.org) to identify the interrelationship of subclustered cells and assign a trajectory based on similarities in expression patterns of each cell. The trajectory infers a transcriptional level relationship of dynamic cellular processes (Saelens et al., 2019). Cells assigned to subclusters served as input which was then ordered into a minimum spanning tree. Paths through the tree are fitted by simultaneous principal curves with each cell’s pseudotime value determined by its projection onto the curves. Slingshot analysis indicated that Ly6C^Hi^ subclusters shift from subcluster 3 (as the root) towards subcluster 0 and 1, ending at subcluster 2 (**Fig. 3C top**). Ly6C^Hi^ subcluster 3 had a prominent proliferative signature marked by expression of *Mki67*, anti-apoptotic gene, *Birc5*, and DNA and nucleosome structural genes (*Top2a, Hist1h1b, H2afz*) (**Fig. 3B bottom**). Ly6C^Hi^ subcluster 0 had a pro-inflammatory signature marked by expression of interferon response genes (*Ifitm3, Lyz, Ifi27l2a, Ccl6, Cxcl2*, and *Il1b*) (**Fig. 3B** and **3C, heatmaps)**. Notably, in the aged mouse brain, there was a significant increase of subcluster 1 cells and subcluster 2 cells. Subcluster 2 appeared to be unique to the aged brain with a higher expression of *MPO*, which is associated with Alzheimer’s disease (Gellhaar et al., 2017; Giri et al., 2017). Ly6C^Hi^ subcluster 1 had enriched expression for MHC-II complex pathways while Ly6C^Hi^ subcluster 2 was enriched for both MHC-II and neutrophil-mediated cytotoxicity (**Fig. 3D** and **Supplementary Table 4**). Enrichment for these pathways indicated that Ly6C^Hi^ monocytes may experience transcriptional plasticity in response to an inflammatory environment in the aged CNS. We also performed re-clustering and subcluster proportional analysis in young and aged mice compared to their ABX treated counterparts. However, we observed minimal shifting of the Ly6C^Hi^ population in aged and young mice experiencing gut dysbiosis (**Supplementary Fig. 4B**), suggesting that the Ly6C^Hi^ plasticity is more greatly affected by age-dependent changes.

**Figure 3:**
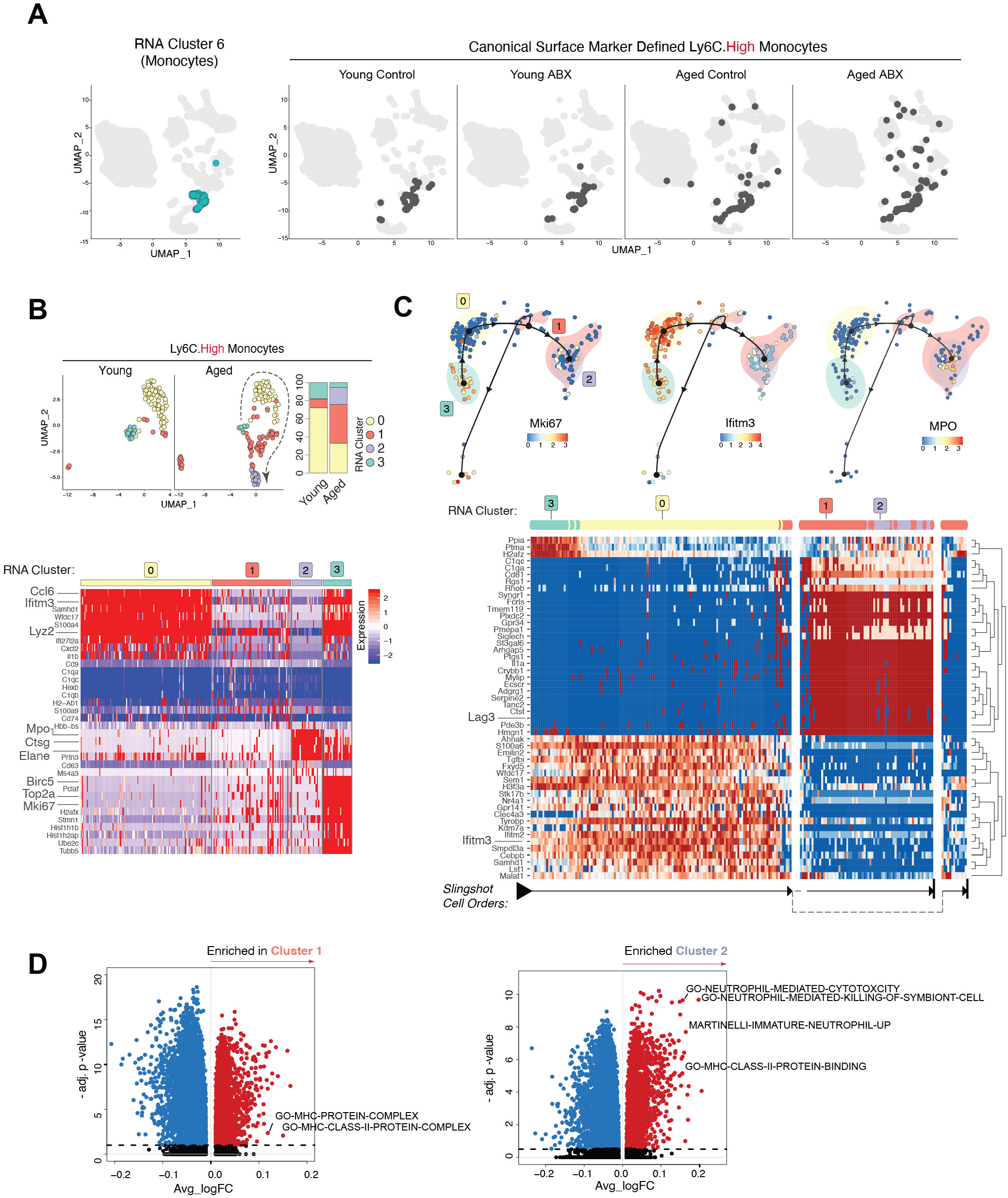
Innate Ly6C^Hi^ monocytes show microenvironmental-dependent plasticity in aged brain. A) (left) UMAP clustering as in (1B) with RNA cluster 6 only in turquoise. (right - 4 plots) UMAP clustering as in (1B) with Cell-ID Ly6C^Hi^ darkened. UMAP plots are split by the sample group. B) (top left) Reclustered Ly6C^Hi^ cells from young and aged mice projected onto UMAP. Plots split by young and aged. (top right) Stacked bar charts of Ly6C^Hi^ subclusters compared between young and aged mice. (bottom) Heatmap of Ly6C^Hi^ subcluster marker genes. C) (top) Slingshot Dynamo ordering of Ly6C^Hi^ subclusters. Plots split by subcluster marker gene expression (*Mki67, Ifitm3*, and *Ctsg*). (bottom) Heatmap of marker genes generated by the Slingshot Dynamo cluster ordering. Arrows at the bottom of the heatmap denote trajectory assignment. D) (left) Volcano plot showing differentially enriched gene pathways in Ly6C^Hi^ subcluster 1 compared to all other Ly6C^Hi^ subclusters. (right) Volcano plot showing differentially enriched gene pathways in Ly6C^Hi^ subcluster 2 compared to all other Ly6C^Hi^ subclusters.

### Aged brain increases Ly6C^Low^ patrolling monocyte plasticity

Next, similar to Ly6C^Hi^ monocytes, we explored the plastic nature of Cx3cr1^High^CCR2^-^Ly6C^Low^ (Ly6C^Low^) monocytes, which are an anti-inflammatory subset of the immune population that patrol within the lumen of blood vessels and promote tissue repair (Auffray et al., 2009). In relation to neurodegenerative disease, Ly6C^Low^ monocytes are reported to be recruited to inflammation in the brain by vascular amyloid-beta (Aβ) microaggregates (Thériault et al., 2015). Upon recruitment, Ly6C^Low^ monocytes perform tissue repair by internalizing and transporting Aβ microaggregates from the brain tissue into circulating blood (Michaud et al., 2013). As natural eliminators of Aβ from brain tissue, Ly6C^Low^ immune cells serve an important role in maintaining brain tissue integrity and in patients with Alzheimer’s disease (Saresella et al., 2014).

The Cell-ID Ly6C^Low^ population exhibited a shifted transcriptional program associated with aging. At the RNA transcriptome level, Ly6C+ monocytes were most abundant in RNA cluster 3, marked by expression of MHC-II genes (*Cd74, H2-Ab1*, and *H2-Aa), Lyz2*, and *Cxcl2* (**Fig. 1C** and **Fig. 4A left**). Visualization of Cell-ID Ly6C^Low^ cells on the transcriptome-based UMAP revealed that these cells spread into varying transcriptional signatures and dispersion was increased in both the aged control and aged ABX conditions (**Fig. 4A right**). Reclustering of these cells revealed six distinct subclusters (**Fig. 4B top left**). Only the aged brain had Ly6C^Low^ cells with the subcluster 5 signature and had a higher abundance of subcluster 1 (**Fig. 4B top right**). Young and aged mice with gut dysbiosis did not exhibit significant shifting in Ly6C^Low^ subclusters in comparison to their counterparts with intact microbiomes (**Supplementary Fig. 5A**).

**Figure 4:**
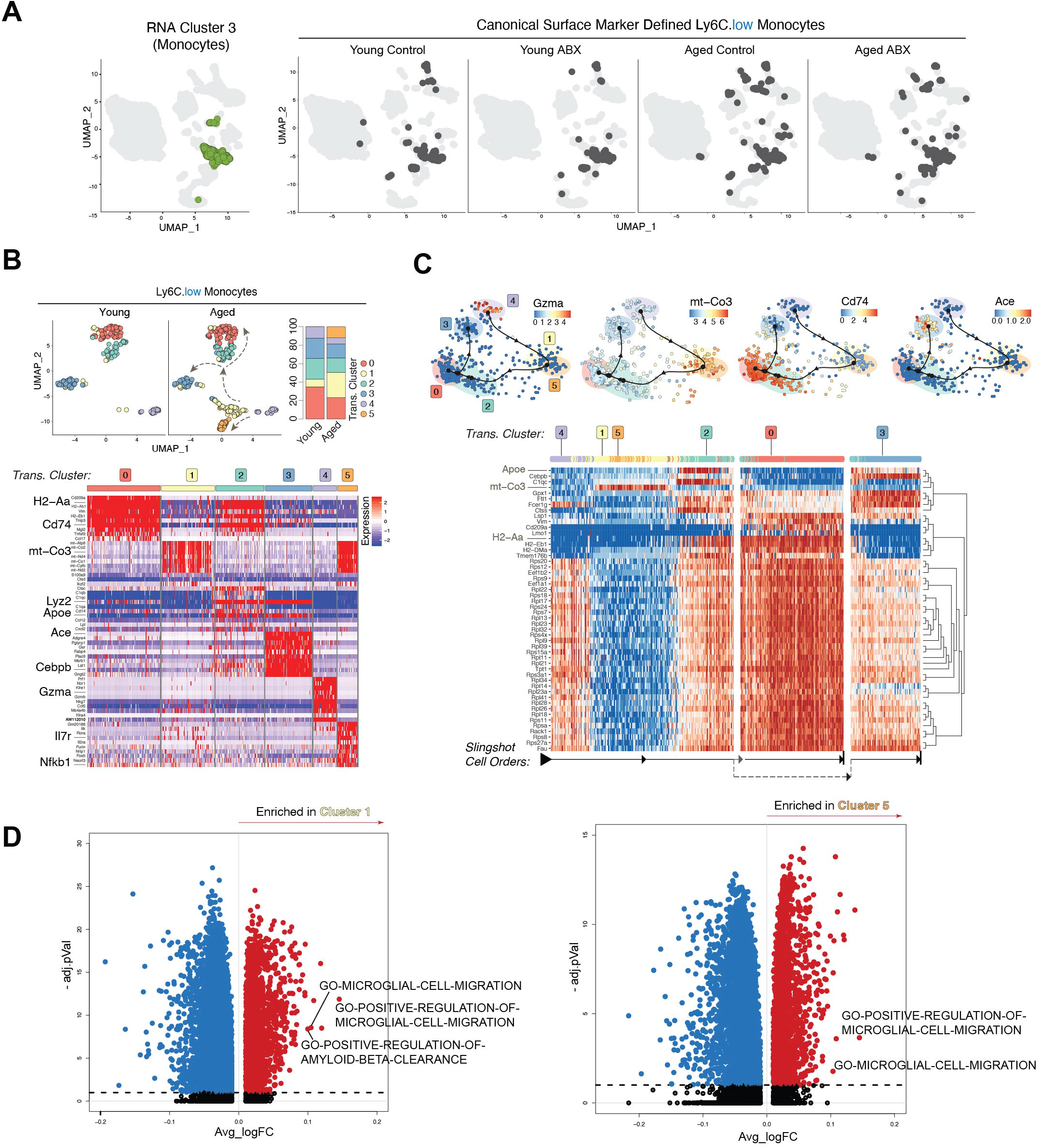
Aged brain increases Ly6C^Low^ patrolling monocyte plasticity. A) (left) UMAP clustering as in (1B) with RNA cluster 3 indicated in green. (right - 4 plots) UMAP clustering as in (1B) with Cell-ID Ly6C^Low^ darkened. UMAP plots are split by the sample group. B) (top left) Reclustered Ly6C^Low^ cells from young and aged mice projected onto UMAP. Plots split by young and aged. (top right) Stacked bar charts of Ly6C^Low^ subclusters compared between young and aged mice. (bottom) Heatmap of Ly6C^Low^ subcluster marker genes generated in Seurat analysis. C) (top) Slingshot Dynamo ordering of Ly6C^Low^ subclusters. Plots split by subcluster marker gene expression (*Gzma, mt-Co3, Cd74* and *Ace*). (bottom) Heatmap of marker genes generated by Slingshot Dynamo cluster ordering. Arrows at the bottom of the heatmap denote trajectory assignment. D) (left) Volcano plot showing differentially enriched gene pathways in Ly6C^Low^ subcluster 1 compared to all other Ly6C^Low^ subclusters. (right) Volcano plot showing differentially enriched gene pathways in Ly6C^Low^ subcluster 5 compared to all other Ly6C^Low^ subclusters.

Both subclusters 1 and 5 had overlapping marker genes and shared high levels of mitochondrial gene expression (*mt-Atp6, mt-Co2*, and *mt-Co3*) (**Fig. 4B bottom** and **Supplementary Table 2**). Trajectory inference analysis and RNA velocity analysis visualized the projected ordering of the Ly6C^Low^ subclusters in which the cells progressively shift in transcriptional signature to the next cellular status **(Fig. 4C** and **Supplementary Fig. 5B)**. Dyno analysis predicted subcluster 4 as the root, marked by high levels of cytotoxic genes (*Gzma* and *Gzmb*), and these cells shifted into closely related subclusters 1 and 5 (**Fig. 4B bottom** and **Fig. 4C top**). With highly expressed mitochondrial genes (e.g. *mt-Co3*), subclusters 1 and 5 seemingly serve as a transitory state before further evolving into subcluster 2, wherein the cells then branch into either subclusters 0 or 3 (**Fig. 4C**). Ly6C^Low^ subcluster 2 had overlapping marker genes of subclusters 0 and 3 with a high expression of *Cd74* and *Lyz2*, suggesting that this cluster serves as a transition state between differentiation into two branches of either subcluster 0 or 3 gene expression programs (**Fig. 4B bottom**). Subcluster 0 was marked by expression of MHC-II genes (*H2-Aa, H2-Ab1*, and *Cd74*), suggesting that these cells upregulated antigen presentation possibly due to increased phagocytosis in response to tissue damage (**Fig. 4B bottom**). Subcluster 3 showed a differentiated cell genotype with high *Ace* and *Cebpb* expression (**Fig. 4B bottom** and **4C top**). *Cebpb* encodes a transcription factor, C/EBPβ, which is reported to drive differentiation of peripheral Ly6C^+^ monocytes into the Ly6C^Low^ monocyte subtype (Mildner et al., 2017). *Ace* encodes angiotensin-converting enzyme (ACE), a peptidase that facilitates myeloid maturation and inhibits the development of myeloid-derived suppressor cells (Shen et al., 2014). Marked expression of *Cebpb* and *Ace* in subcluster 3 suggests that these Ly6C^Low^ cells are actively differentiating into mature monocytes. The increased proportion of subclusters 1 and 5 in the aged brain suggests that Ly6C+ cells have an increased active differentiation and response to the microenvironmental changes. LPS stimulation has reportedly induced *IL7r* expression in monocytes to regulate T cell homeostasis (Al-Mossawi et al., 2019). Distinct from subcluster 1, subcluster 5 had high expression of *Il7r* and *Nfkb1*, suggesting that these cells are potentially responding to increased neuroinflammation in the aged brain and could be involved in regulating T cell response (**Fig. 4B bottom**). Both subclusters 1 and 5 were enriched for pathways involved in microglia cell migration, which may be indicative of more general macrophage migration toward sites of injury and response to inflammatory stimuli (**Fig. 4D** and **Supplementary Table 4**). Notably, subcluster 1 was also enriched for gene pathways in amyloid-beta clearance (**Fig. 4D left**), which have been observed in CNS phagocytic cells in response to neuroinflammation and Alzheimer’s disease (Saresella et al., 2014; Thériault et al., 2015).

### CNS innate lymphoid cell (ILC) plasticity reflects chronic neuroinflammation in the aged brain

ILCs originate from the same lymphoid progenitor as B and T lymphocytes but lack antigen-specific receptors and play important regulatory functions in organ-specific immunity (Colonna, 2018). The plasticity among ILC subtypes in peripheral tissues has become increasingly appreciated (Bal et al., 2020; Colonna, 2018) and recent studies have started to demonstrate the role of ILCs in CNS diseases, such as neuroinflammation (Kwong et al., 2017; Romero-Suárez et al., 2019). However, how ILCs respond to systemic factors, such as aging and gut dysbiosis, have not been thoroughly investigated. To begin delineating the plastic nature of ILCs derived from CNS tissue, we distinguished Cell-ID ILCs (ILC-like cells) from T cells by CD117 surface expression (CD117+) and low CD3 surface expression (**Supplementary Fig. 2, ILC Gating**). ILC-like cells made up 2.5% of the total leukocytes we sequenced and predominantly exhibited transcriptome signatures of RNA-based clusters 7 and 9. (**Fig. 5A left** and **Supplementary Table 3**). Similar to the Ly6C+ monocytes, we observed increased dispersion of ILC-like cells into several RNA-based clusters (**Fig. 5A right**). Interestingly, there was a significant dispersion of ILC-like cells from the aged control group into RNA cluster 3, primarily representing a CD8+ T cell-like transcriptome signature. This was not observed in the aged ABX group indicating that gut microbiota-immune signaling may mediate ILC transcriptome plasticity within the aged brain.

**Figure 5:**
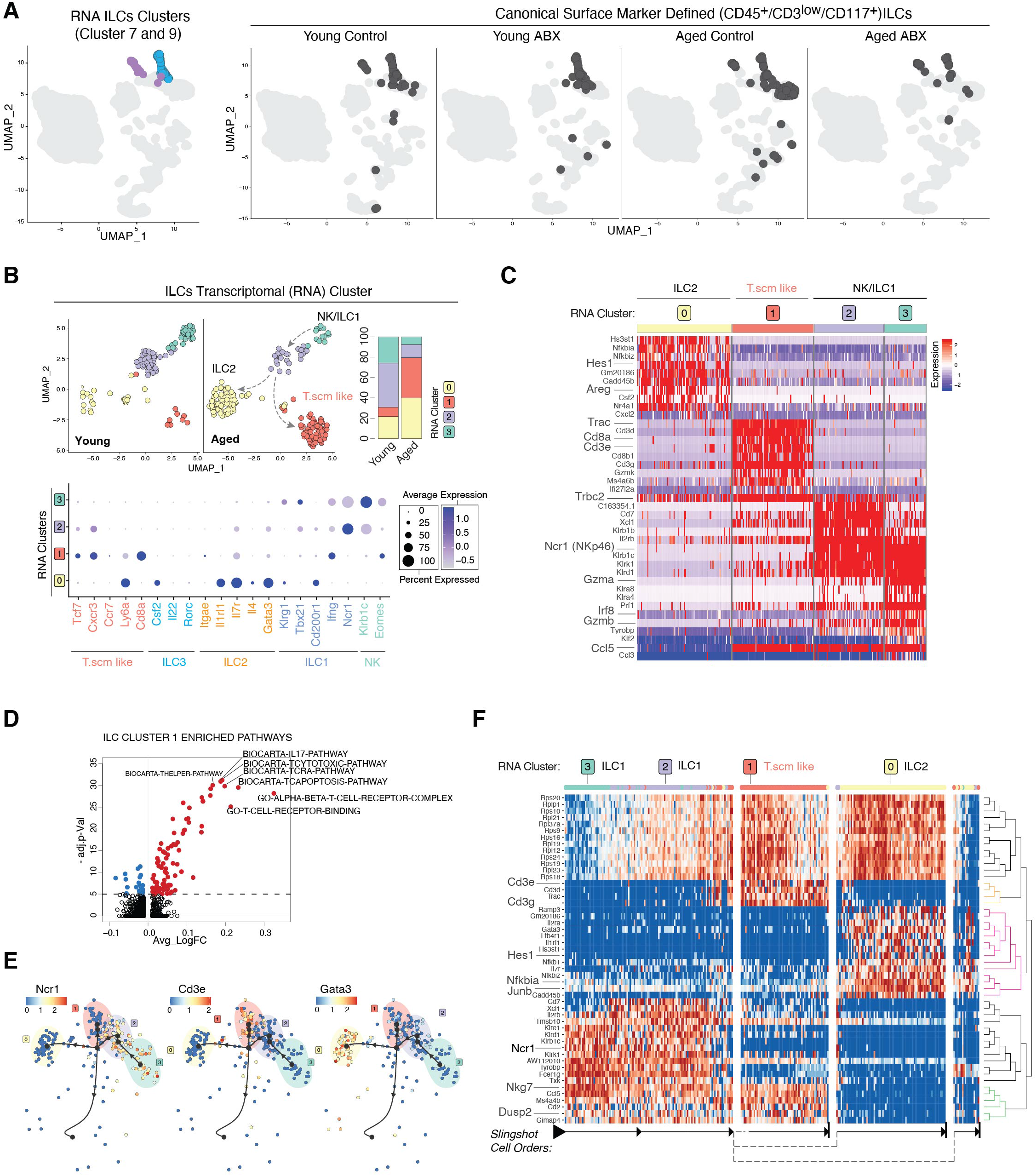
CNS innate lymphoid cell (ILC) plasticity reflects chronic neuroinflammation in the aged brain. A) (left) UMAP clustering as in (1B) with RNA-based clusters 7 (blue) and 9 (purple) highlighted. (right - 4 plots) UMAP clustering as in (1B) with Cell-ID ILCs darkened. UMAP plots are split by the sample group. B) (top left) Reclustered ILCs from young and aged mice projected onto UMAP. Plots split by young and aged. (top right) Stacked bar charts of ILC subclusters compared between young and aged mice. (bottom) Dot plot showing ILC subcluster expression level and percentage of ILC subtype, T memory stem cell (T_scm_) and Natural Killer (NK) marker genes. The color scale represents the level of expression and the size of the dot represents the percent of the population expressed in the indicated gene. C) Heatmap of ILC subcluster marker genes. D) Volcano plot showing differentially enriched gene pathways in ILC subcluster 1 compared to all other ILC subclusters. E) Slingshot Dynamo ordering of ILC subclusters. Plots split by subcluster marker gene expression (*Ncr1, Cd3e*, and *Gata3*). F) Heatmap of marker genes generated by Slingshot Dynamo cluster ordering. Arrows at the bottom of the heatmap denote trajectory assignment.

To further discern the nature of ILCs plasticity, we subsetted the ILC-like cells and reclustered the cells based on their single-cell transcriptome (**Fig. 5B)**. We first focused on ILC-like cells from young and aged brains and found that they segregated into four transcriptionally distinct clusters (**Fig. 5B top left**). Aged ILC-like cells had significantly higher proportions of subclusters 0 and 1 (**Fig. 5B top right**). Using previously described ILC subtype marker genes, we identified ILC and lymphoid subtypes present among the ILC-like subclusters (Bal et al., 2020; Colonna, 2018; Gury-BenAri et al., 2016). Subcluster 0 was marked by a higher expression of *Gata3, Il1rl1, Itgae*, and *Areg*, resembling the ILC2 subtype (**Fig. 5B bottom, Fig. 5C** and **Supplementary Table 2**). Interestingly, subcluster 1 cells did not express previously defined ILC subtype marker genes. Instead, they showed a T cell-like signature with expressions of *Cd3e, Cd8a, Tcf7*, and *Trac* (**Fig. 5B bottom, Fig. 5C** and **Supplementary Table 2**). Of note, this T cell-like subcluster had a significantly lower CD3 surface expression than canonical T cells (**Supplementary Fig. 2, ILC gating**), and negative to low expression levels of *Cd8a* and *Cd3e*, further confirmed at the RNA level (**Supplementary Fig. 7A**). We also noted that most of *Cd3e+* or *Cd5+* cells did not express *Tbx21* (T-bet), an ILC marker gene (**Supplementary Fig. 6A**) (Colonna, 2018). Instead, the ILC T cell-like subcluster expressed marker genes observed in T memory stem cells (T_scm_) including *Tcf7, Cxcr3* and *Ly6a* (Gattinoni et al., 2017), T memory cell transcription factor *Eomes* (**Fig. 5B bottom** and **Supplementary Table 2)**, and brain-resident memory T cell marker *Ccl5* (Knox et al., 2014; Steinbach et al., 2019) (**Supplementary Fig. 6B)**. These T_scm_-like cells were also enriched with T helper, IL-17 and T-cytotoxic pathways (**Fig. 5D** and **Supplementary Table 4**). Subclusters 2 and 3 had some overlapping expression of NK-like genes (*Klrb1c, Klrk1* and *Ncr1*) (**Fig. 5C** and **Supplementary Table 2**). However, a higher expression of T-bet transcription factor, *Tbx21*, in subcluster 2 indicates this cluster resembled ILC1 status whereas expression of *Eomes* in subcluster 3 designates an NK-like status (Bal et al., 2020). There were no cells expressing the ILC3 marker gene *Rorc* (**Fig. 5B bottom**). Next, we used RNA velocity analysis (**Supplementary Fig. 6C)** and slingshot trajectory analysis to order the cellular states and assign a potential inter-lineage transition path of the ILC-like subclusters. Based on cell velocity analysis and Slingshot trajectory, subcluster 3 with the NK-like transcriptome was assigned as the cell type of origin and appears to shift into the closely related subcluster 2 (ILC1). From there, differentiation was projected to split into subcluster 1 with T_scm_-like transcriptome and subcluster 0 (ILC2) (**Fig. 5E-F**). The trajectory projections indicated potential transdifferentiation of NK-like cells into ILC1s which then branched into either ILC2 or brain resident T_scm_-like in the aged brain tissue environment (**Fig. 5B** indicated by arrows). The enrichment of ILC2 and T_scm_-like cells suggests an increased prevalence of type 2/chronic inflammatory response occurring in the aged brain tissue environment (Steinbach et al., 2019; Zaiss et al., 2015).

### Antibiotics treatment alters ILC plasticity pattern in the aged mice, but not in the young mice

Peripheral ILCs are known to have the ability to recognize both self and pathogenic molecules and respond to systemic cues, e.g. gut microbiota dysbiosis (Bal et al., 2020; Colonna, 2018; Gury-BenAri et al., 2016). Upon gut microbiota dysbiosis, intestinal ILCs underwent extensive epigenetic and transcriptional changes, which led to the expansion of ILC3s in either germ-free or ABX treated mice (Gury-BenAri et al., 2016). This finding prompted us to further investigate the impact of ABX treatment on CNS-derived ILC transcriptome plasticity. We reclustered ILC-like cells from the young control vs. young ABX and aged control vs. aged ABX groupings. For both young and aged groups, the ILC-like cells separated into three transcriptionally distinct clusters (**Fig. 6A, left**). Interestingly, ABX treatment led to a significant decrease in the frequency of subcluster 1 only in aged mice, while subcluster distributions in young mice with gut dysbiosis remained relatively stable (**Fig. 6A right**). Subcluster 2 resembled the ILC1 subtype signature (*Ncr1, Tbx21*, and *Gzma*) while subcluster 0 resembled the ILC2 subtype with the expression of *Hes1, Areg*, and *Gata3* (**Fig. 6B** and **Supplementary Table 2**). Subcluster 1 closely mirrored the T_scm_-like cluster we previously described in the aged control mice, with the expression of *Ly6a, Cd8a*, and *Cxcr3* (**Fig. 6B** and **Supplementary Table 2**). Slingshot Dyno analysis ordered the subclusters and projected differentiation similarly to ILC-like cells compared between young and aged mice, with the ILC1 subcluster projected to differentiate and split into the T_scm_-like cells and ILC2s (**Fig. 6C-D).** Of note, while ABX treatment prevented the transdifferentiation path from ILC1 to T_scm_-like cells in the aged brain microenvironment, ABX treatment had no impact on classic Cell-ID CD8+ T cells at both compositional and transcriptional levels (**Supplementary Fig. 7B-D**), highlighting the unique identity of T_scm_-like cells in the aged brain. While potential transdifferentiation between ILC1 and T_scm_-like cells in the aged brain requires further functional validation, our data provide the first evidence that ILC plasticity associated with gut microbiota depletion is not restricted to intestinal ILCs but affects ILCs found in the brain as well.

**Figure 6:**
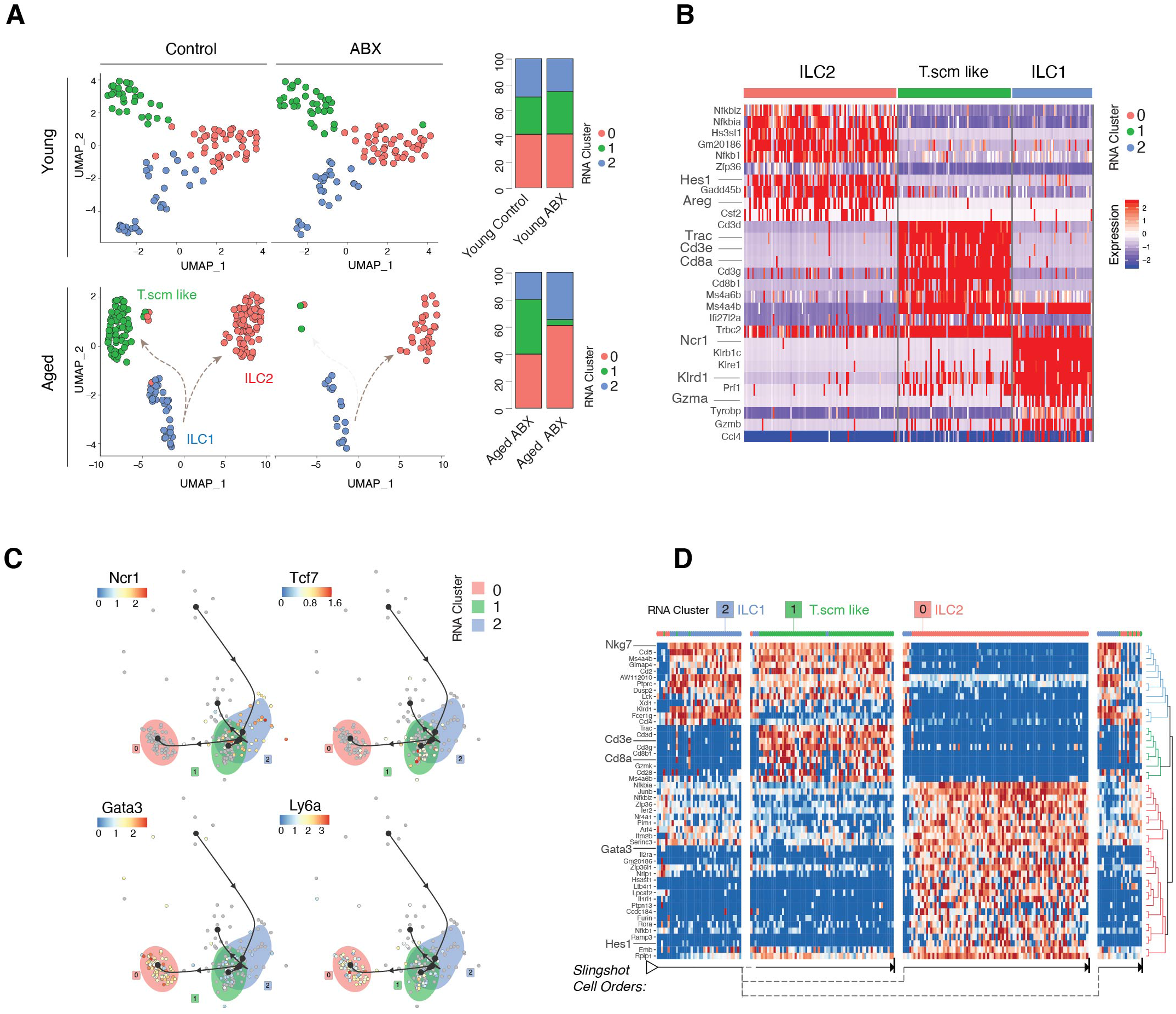
Antibiotics treatment alters ILC plasticity pattern in the aged mice. A) (top left) Reclustered ILCs from young control and young ABX mice projected onto UMAP. Plots split by young control and young ABX. (top right) Stacked bar charts of ILC subclusters compared between young control and young ABX mice. (bottom left) Reclustered ILCs from aged control and aged ABX mice projected onto UMAP. Plots split by aged control and aged ABX. (top right) Stacked bar chart of ILC subclusters compared between aged control and aged ABX mice. B) Heatmap of aged control and aged ABX ILC subcluster marker genes. C) Slingshot Dynamo ordering of aged control and aged ABX ILC subclusters. Plots split by subcluster marker gene expression (*Ncr1, Tcf7, Gata3*, and *Ly6a*). D) Heatmap of marker genes generated by Slingshot Dynamo cluster ordering. Arrows at the bottom of the heatmap denote trajectory assignment.

## DISCUSSION

Defining the complex CNS immune landscape had long been an uncharted area of research until recent advances in single-cell technologies. The canonical identity of immune cells is traditionally defined by their surface protein expression, whereas the functional status of a given immune cell in response to stimuli is often best defined by single-cell transcriptome analysis (Papalexi and Satija, 2018). Previous studies used either CyTOF or scRNA-seq to characterize CNS immune cells (Jordão et al., 2019; Mrdjen et al., 2018; Ximerakis et al., 2019). Due to the technical nature of CyTOF and scRNA-seq, neither technique can capture a holistic image of both canonically-defined lineage identity and detailed functional status at the transcriptome level. Depending on the experimental context, the observed surface marker overlap, as well as shared transcriptome signatures, often led to debates on the meaningful definitions of immune cell subsets (Ealey and Koyasu, 2017; Simoni and Newell, 2017). To address this challenge, CITE-seq was developed as the ideal multi-modal sequencing tool to track transcriptional changes happening within a specific cell type by directly coupling surface marker expression profiling with single-cell transcriptome profiling (Stoeckius et al., 2017). In this study, enabled by multi-modal CITE-seq analysis, we attempted to examine the transcriptional plasticity of canonically identified immune cells in the brain. We systematically delineated compositional and transcriptional changes of the brain immune landscape in response to the systemic influence of aging and gut dysbiosis. In particular, our results highlighted the highly plastic nature of monocyte subsets and innate lymphoid cells in response to systemic changes, suggesting their potential role in the aged brain and gut-brain communication.

Microglia and BAMs are the most well studied CNS-resident innate immune cells. Using CyTOF analysis, Mrdjen *et al.* described changes in surface expression level and compositional changes among the CNS resident immune populations in response to aging and neuroinflammation (Mrdjen et al., 2018). In the brain of geriatric mice, they reported an increased frequency of MHCII+CCR2-BAMs. Interestingly, a subset of microglia expressed a higher level of phagocytosis-associated surface markers, CD11c, in geriatric mice (Mrdjen et al., 2018). Consistent with the above observations, our study provided further single-cell transcriptome insights on aging-induced changes (**Fig. 2**). Although the transcriptome of microglia and BAMs were relatively stable and less plastic compared to monocytes and ILCs, it was evident that these cells were shifted toward a pro-inflammatory and antigen presentation program (RNA cluster 1 in microglia and RNA cluster 3 in BAMs) under the influence of aging, and marginally so by gut dysbiosis (**Fig. 2**). We suspect that such shifting of microglia and BAMs with a pro-inflammatory profile is reflective of a more inflamed environment within the aged brain (Erickson and Banks, 2019; Plaza-Zabala et al., 2017). Despite previous evidence of a dysfunctional microglia immune response in germ-free (GF) mice, it is worth noting that we observed that ABX-induced gut dysbiosis only marginally impacted microglia and BAM homeostasis, which is consistent with a previous report (Erny et al., 2015). This observation suggests that under normal physiological conditions, the CNS resident myeloid cell compartment remains largely resistant to the gut dysbiosis induced changes.

In contrast to microglia and BAMs, the peripheral myeloid cells, such as Ly6C+ monocytes and macrophages, have been known to demonstrate considerable functional plasticity in response to infection, injury, or disease (Galli et al., 2011). We observed that peripherally-derived monocytes showed substantial plasticity under the influence of systemic changes, mostly associated with aging, but not ABX-induced gut dysbiosis. In the aged brain, Ly6C^Hi^ inflammatory monocytes had an increased inflammatory status and dispersion into neighboring clusters (**Fig. 3**) and Ly6C^Low^ patrolling monocytes has enhanced migratory gene signatures (**Fig. 4**), suggesting their roles in guarding brain homeostasis.

Differing from myeloid cells, ILC-like cells displayed substantial transcriptional plasticity in response to both aging and gut dysbiosis stimuli (**Fig. 5-6**). It has been increasingly appreciated that peripheral ILCs substantially mirror the T helper cell signature and have been regarded as important mediators in inducing and resolving inflammation as well as maintaining tissue homeostasis and mucosal barriers (Bal et al., 2020; Colonna, 2018; Derecki et al., 2019). Compared with peripheral ILCs, the ILCs derived from brain tissue have not been extensively studied. Relative to ILC-like cells of young mice, ILC-like cells in aged mice shifted their transcriptional program from the ILC1 into ILC2 and T_.scm_-like subtypes (**Fig. 5B, E, and F**). The ILC2 subtype is marked by a high expression of amphiregulin (AREG) (Colonna, 2018) (**Fig. 5C**). AREG is reported to be an epidermal growth factor (EGF)-like molecule that may be critical in Type-2 mediated resistance and tolerance and can be expressed by multiple immune populations in a variety of inflammatory conditions (Zaiss et al., 2015). Importantly, Zaiss *et al.* and others have reported that AREG expression appears to be a key factor in inducing tolerance in order to maintain tissue integrity following damage or in chronically inflamed environments (Zaiss et al., 2015). The increased disease-associated BAMs and inflammatory expression in Ly6C+ monocytes (reported in **Fig. 2-4**) in our data demonstrate that age-associated neuroinflammation is occurring. The shifting of ILC-like cells from ILC1 to ILC2 (AREG+) (**Fig. 5B**) supports the notion that ILCs change their transcriptional signature in response to chronic neuroinflammation to maintain brain tissue integrity. In addition, recent studies on CNS ILCs revealed essential roles of ILCs in guiding T cell CNS infiltration as well as T cell engagement and response to neuroinflammation (Kwong et al., 2017). Thus, the observed shifting to ILC2 subtypes within the aged brain potentially regulates the CD8+ T cell infiltration in the aged brain observed in our study (**Fig. 1D-F**) as well as in recent reports (Dulken et al., 2019; Mrdjen et al., 2018).

Most interestingly, ABX treatment specifically reduced ILC-like subclustered cells with the T_scm_-like transcriptome signature only in the aged brain, but not in young mice (**Fig. 6A**). These T_scm_-like cells were identified based on the surface protein expression pattern of CD117+/CD3^low^/CD8^low^, yet express many T cell-like transcriptome qualities. Since CD8+ T cell infiltration was maintained in the aged ABX mice with reduced T_scm_ cell frequency (**Supplementary Fig 7A**), these T_scm_-like cells are not necessarily required for CD8+ T cell infiltration in the aged brain. In addition, ABX treatment sensitive T_scm_-like cells appear to function independently to the classic CD8+ T cells, which were resistant to ABX-treatment-induced changes at the transcriptional level (**Supplementary Fig. 7B-D**). To our knowledge, our study provides one of the first accounts of gut dysbiosis affecting a specific subset of brain-derived T_scm_-like ILCs. The functional significance of the T_scm_-like ILCs in the aging brain and how their plasticity is influenced by dysbiosis in the aged tissue environment warrants further research.

In summary, our study provides a holistic map of the brain immune plasticity in response to aging and gut dysbiosis. The molecular characterization of the influential impact of aging on both innate and adaptive immune systems in the brain will guide further mechanistic studies on age-associated neuroinflammation and other neurological diseases.

## ACKNOWLEDGMENTS

This work was partially funded by NIH R01 CA194697-01 (S.Z.), NIH R01 CA222405 - 01A1 (S.Z.) and the University of Notre Dame CRND Catalyst Award (S.Z.). We would like to acknowledge and thank the Dee Family endowment (S.Z.). We are additionally grateful for the technical support of the following core facilities: Notre Dame Genomics and Bioinformatics Core Facility, Notre Dame Freimann Life Sciences Center, Indiana University School of Medicine Center for Medical Genomics and Indiana University Simon Cancer Center CTSI Center for Medical Genomics Core Facility.

## AUTHOR CONTRIBUTIONS

S.M.G., I.H.G., and S.Z. conceived the original hypothesis and designed experiments.

S.M.G., I.H.G., Q.W., B.P., J.L., and S.Z. performed experiments.

S.M.G., I.H.G., A.Z., and S.Z. analyzed data.

K.Y. contributed critical intellectual guidance to this study.

S.M.G., I.H.G., and S.Z. wrote and revised the manuscript.

S.Z. supervised the study.

All authors reviewed the manuscript.

## DECLARATION OF INTERESTS

The authors declare no competing interests.

## METHODS

### Mice

8-week old and 68-week old SPF C57BL/6 mice were purchased from Jackson Laboratory. Mice were acclimated in the animal facility for 2 weeks and randomized within age groups to normalize the gut microbiota. 10-week and 70-week old mice were assigned to either vehicle or antibiotics (ABX) treatment groups. The ABX groups received an ABX cocktail containing metronidazole (MedChem Express, 0.25 g/L), vancomycin hydrochloride (BioVision, 1 g/L), neomycin (VWR, 1 g/L), and ampicillin sodium salt (Sigma, 0.5 g/L). Initially, the ABX was delivered in the drinking water and by oral gavage every other day to all mice for the first week of treatment. Then, all mice in the study were switched to the vehicle drinking water and received oral gavage of either vehicle or ABX cocktail twice daily for an additional 2 weeks of treatment. Mice were weighed every 3-4 days during treatment to check that healthy body weight was maintained. Fresh ABX cocktail was prepared every 3 days for oral gavage. Water bottles were refreshed every 3-4 days to ensure mice received active antibiotics each dose. Mouse fecal samples were collected on day 0 and day 21 (endpoint) of treatment for 16S analysis. After ABX pretreatment was completed, mice were sacrificed to collect brain tissue for 10X Genomics Chromium Single Cell Gene Expression analysis. All animal studies were performed ethically and in accordance with the IACUC protocol approved by the University of Notre Dame IACUC committee.

### 16S rRNA sequencing and analysis

Mouse fecal DNA extraction was performed using the ZymoBIOMICS DNA Miniprep Kit (Cat# D4300). 16S V3/V4 region amplification and library preparation were performed following the Illumina 16S Metagenomic Sequencing Library Preparation (PN 15044223 Rev. B). Libraries were submitted to the University of Notre Dame Genomics & Bioinformatics Core Facility (GBCF) for Illumina sequencing. GBCF normalized and multiplexed the libraries into a single pool prior to validation by Qubit High-Sensitivity dsDNA, Agilent Bioanalyzer DNA 7500 Chip, and Kapa Illumina Library Quantification qPCR analyses. The final library (8.5pM) and PhiX (0.7pM) were sequenced on the MiSeq System using MiSeq V2 500 cycle kit (250-PE with 8-bp dual indexing). Base calling and demultiplexing were performed by MiSeq Controller Software. After quality control filtering; a total of 2,734,528 reads were processed with an average of 171,159 reads per sample. We used the Divisive Amplicon Denoising Algorithm version 2 (DADA2) R package to analyze the 16S amplicon reads and construct the operational taxonomic units (OTUs). This package incorporates a quality-aware model of Illumina amplicon errors to improve the identification of real variants and minimize false positives (Callahan et al., 2016).

### Tissue collection and single cell preparation

For CITE-seq brain tissue collection, mice were anesthetized with isoflurane and transcardially perfused with cold 1x PBS. Cells were isolated from mouse brain tissue by digesting the brains into single cell suspensions and then enriched from other neural cells by density gradient centrifugation as follows: Brains from mice transcardially perfused with 1x PBS were extracted, minced with scissors and triturated with a P1000 micropipette. The resulting brain tissue slurry was centrifuged at 300g for 2 minutes. The supernatant was removed and the pellet resuspended and processed as directed by the Multi-tissue Dissociation Kit I (Miltenyi Biotec, 130-110-201). Brains were enzymatically digested into a single cell suspension by rotating at 37°C for approximately 25 minutes with trituration halfway through incubation. The resulting cell suspension was strained through a 100μm cell filter as needed, diluted in 1x HBSS, and centrifuged at 300g for 10 minutes. The resulting supernatant was discarded and the pellet resuspended in 3mL 70% Percoll (GE Healthcare, 17-0891-02). The Percoll gradient and density layering were prepared (from bottom to top: 70%, 37%, 30%). Density gradient centrifugation was performed for 20 minutes at 2000rpm with no break. Mononucleated cells were isolated from the buffy layer between the interface of the clear Percoll and red-colored Percoll. Following washing in 1x HBSS, the resultant cell suspension was processed as required for CITE-seq.

### CITE-seq antibodies and cell staining

Following gradient centrifugation, samples were prepared for 10X Genomics Chromium Single Cell Gene Expression analysis as described in the CITE-seq and cell hashing protocol on the CITE-seq website (https://citeseq.files.wordpress.com/2019/02/cite-seq_and_hashing_protocol_190213.pdf). Briefly, samples were blocked by incubation with TruStain fcX in a 50 μL cell staining buffer for 20 minutes on ice. Following the blocking step, samples were stained with Total-seq antibodies purchased from BioLegend: CCR2/CD192 (SA203G11, 150625), CD117/c-kit (2B8, 105843), CD11c (N418, 117355), CD172a/SIRPα (P84, 144033), CD38 (90, 102733), CD44 (IM7, 103045), CD45R/B220 (RA3-6B2, 103263), CD8a (53-6.7, 10073), CD90.1 (OX-7, 202547), Cx3cr1 (SA011F11, 149041), F4/80 (BM8, 123153), I-A/I-E (M5/114.15.2, 107653), Ly6C (HK1.4, 128047), Ly6G (1A8, 127655), NK1.1 (PK136, 108755), PD-1 (RMP1-30, 109123), PD-L1 (MIH6, 153604), CD169/Siglec-1 (3D6.112, 142425), Siglec-H (551, 129615), TMEM119 (A16075D, 853303), XCR1 (Zet, 148227), CD24 (M1/69, 101841), CD103 (2E7, 121437), CD64 (X54-5/7.1, 139325), CD83 (Michel-19, 121519), CD45 (30-F11, 103159), Cd11b (M1/70,101265), CD86 (GL-1, 105047), CD3 (17-A2, 100251), CD4 (RM4-5,100569), and CD25 (PC61, 102055). Additionally, each sample was stained with one unique hashing antibody purchased from BioLegend: HTO1 (M1/42; 30-F11, 155801), HTO2 (M1/42; 30-F11, 155803), HTO3 (M1/42; 30-F11, 155805), HTO4 (M1/42; 30-F11, 155807), HTO5 (M1/42; 30-F11, 155809), HTO6(M1/42; 30-F11, 155811), HTO7(M1/42; 30-F11, 155813), HTO8 (M1/42; 30-F11, 155815). After 25 minutes of staining, samples were washed 4 times prior to delivering the prepared samples to the University of Notre Dame Genomics & Bioinformatics Core Facility (GBCF) for 10X Genomics Chromium Single Cell Gene Expression analysis.

### CITE-seq library preparation and Illumina sequencing

For CITE-seq experiments, GBCF prepared 10X Genomics Chromium Next GEM Single Cell 3’ v3.1 libraries with modification following the CITE-seq protocol Version 2019-02-13 (New York Genome Center Technology Innovation Lab) (https://citeseq.files.wordpress.com/2019/02/cite-seq_and_hashing_protocol_190213.pdf). Libraries (cDNA-, ADT-, and HTO-derived) were validated by Qubit High Sensitivity ds DNA and Agilent Bioanalyzer DNA High-Sensitivity Chip analyses. Afterwhich, libraries were submitted to Indiana University School of Medicine Center for Medical Genomics for multiplexing into a single pool and Illumina sequencing. The multiplex pool contained cDNA library, ADT-derived library, and HTO-derived library by molarity as reported by Agilent TapeStation System to achieve final reads ~50,000-70,000 cDNA reads/cells, ~3,000-5,000 ADT reads/cells and ~2,000 HTO reads/cell. The final library pool was sequenced on the NovaSeq6000 System using NovaSeq XP kit and NovaSeq S2 flow cell (100 cycle) kit (Read1 26-bp, Read2 91-bp and Index1 8-bp). The raw base sequence calls were demultiplexed into sample-specific cDNA, ADT and HTO FASTQ files with bcl2fastq2 Conversion Software v2.20 through CellRanger V3.1.0.

### CITE-seq analysis and statistical analysis

Raw FASTQ files were processed using the Cellranger V3.1.0 software package (10X genomics Inc.) for RNA expression matrix and CITE antibody counts matrix. The data from young and aged generated from two GEM wells were combined using a cellranger aggr pipeline (10X genomics Inc.). Cells were sequenced to comparable sequencing depths (50,084–69,026 reads/cell) and had a similar median unique molecular identifier (UMI) count and median gene number in all conditions. Filtering, normalizing and demultiplexing at the cell and gene levels resulted in a final set of 22,278 cells with 1,128-3,515 cells per sample. The RNA and CITE expression matrices were further analyzed in the Seurat_3.1.1 R package. After pre-filtering and quality control based on minimum gene and cell observance frequency cut-offs, SCTransform function was applied to normalize and scale the data. Dimensionality reduction by principal component analysis and UMAP embedding was performed for cell cluster visualization. Cells were demultiplexed to their original sample groups using the Cell Hashing tags (HTOs). The trajectory inference (TI) analysis was conducted using the Dyno package (https://dynverse.org/) (Saelens et al., 2019). The most appropriate method (Slingshot) was selected based on Dyno recommendations. RNA velocity was performed based on a previous publication (http://velocyto.org/) (Manno et al., 2018). When conducting TI analysis, we also cross-referenced with RNA velocity analysis results to ensure biological meaningful interpretations.

### Gating Strategy

Canonical cell gating was performed as described previously (Mrdjen et al., 2018) and shown in Supplemental Figure 2 based on the CITE-antibody reads, and in instances when a CITE antibody was not available or the reads were not clear, RNA reads for particular markers were used as a substitute and/or a filter to increase confidence. Briefly, CD45 expression level was used to segregate brain-resident (CD45^Low^) and peripherally-derived (CD45^High^) immune cells. CD45^Low^ immune cells were further segregated on the basis of CD38, MHCII, *Tmem119*, and *Mrc1* into microglia (CD38-,MHCII-/Low/*Tmem119*+,*Mrc1*-/Low) or BAM (CD38+,MHCII+*Tmem119*-,*Mrc1+*). Within the CD45^High^ peripheral immune populations, B cells were first identified on the basis of high CD45R-B220 and *Cd19* expression. The remaining CD45^High^ cells were segregated into potential NK and T cell populations or BMDM populations based on moderate to high or low expression of *Thy1* (pan t cell marker) and *Itga2* (highly expressed in T cells and NK cells), respectively. From the *Thy1+Itga2+* population, NK cells were identified on the basis of *Klrb1* positivity, ILCs were identified based on CD3-CD117+, and T cell subsets were identified based on CD3+ and CD4+ or CD8+. The potential BMDM population was filtered to exclude cells with low CD11b and *Itgam* expression and then gated based on Ly6C and Ly6G expression to identify Ly6CLow/NegLy6G-patrolling monocytes, Ly6CHighLy6G-inflammatory monocytes, and Ly6C+Ly6G+ neutrophils. We verified the accuracy of the gating by examining the RNA expression of CNS native myeloid marker genes; *Tmem119*, and *Mrc1* and BMDM-specific genes; *Itga2, Thy1*, and *Itgam* in the identified cell populations.

### Data Availability

CITE-seq data has deposited to GEO with accession number is GSE148127 and is retrievable from GEO with token, mxohuycglzkdzoj, prior to publication, and will be publicly available without the need for a token upon formal acceptance of the manuscript. All other data is available from the corresponding PI upon reasonable request.

### Code Availability

All codes for all CITE-seq analyses are available from the corresponding PI upon reasonable request.

---

**Supplementary Figure 1.**
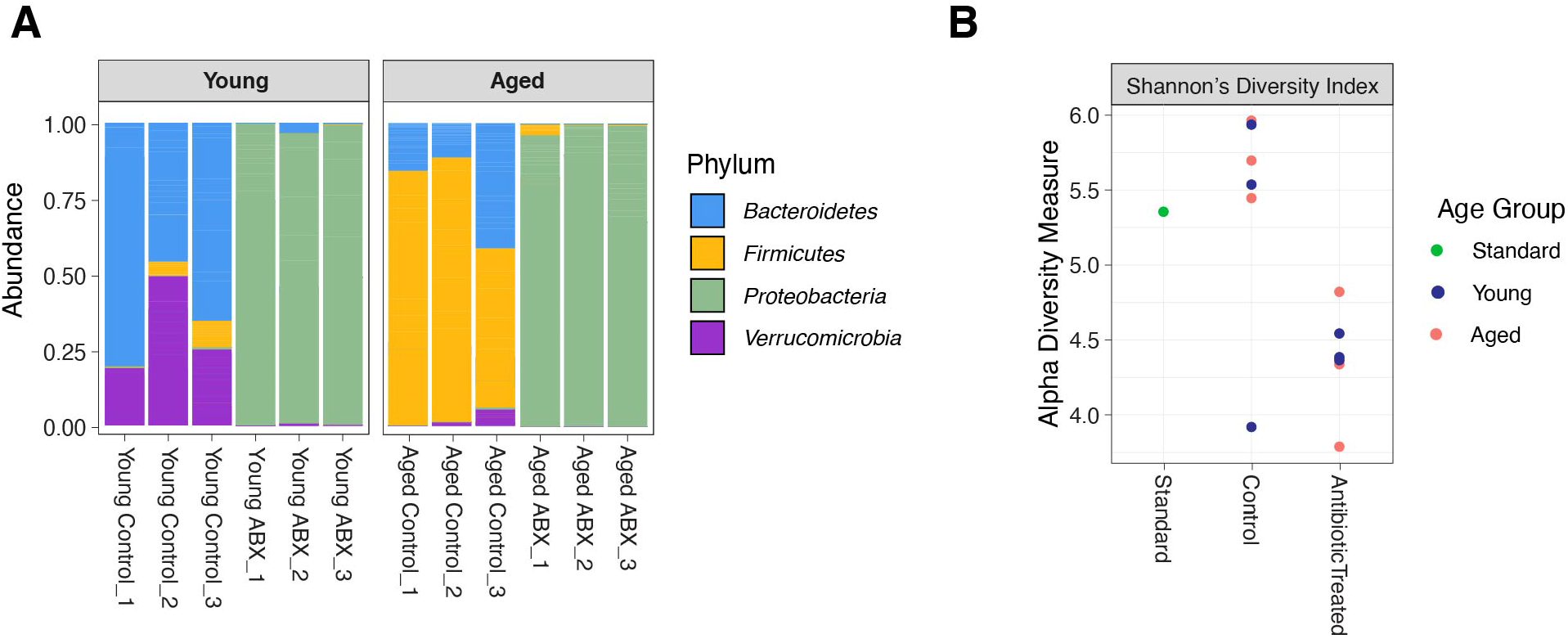
16S analysis of microbiota-depletion efficiency. A) Bacterial phylum abundance frequency measured in young (left) and aged (right) mice receiving vehicle or ABX treatment for 21 days. Three biological repeats for each group. B) Alpha diversity measurement using Shannon’s Diversity index for young and aged mice receiving vehicle or ABX treatment in comparison to standard control. Three biological repeats for each group.

**Supplementary Figure 2.**
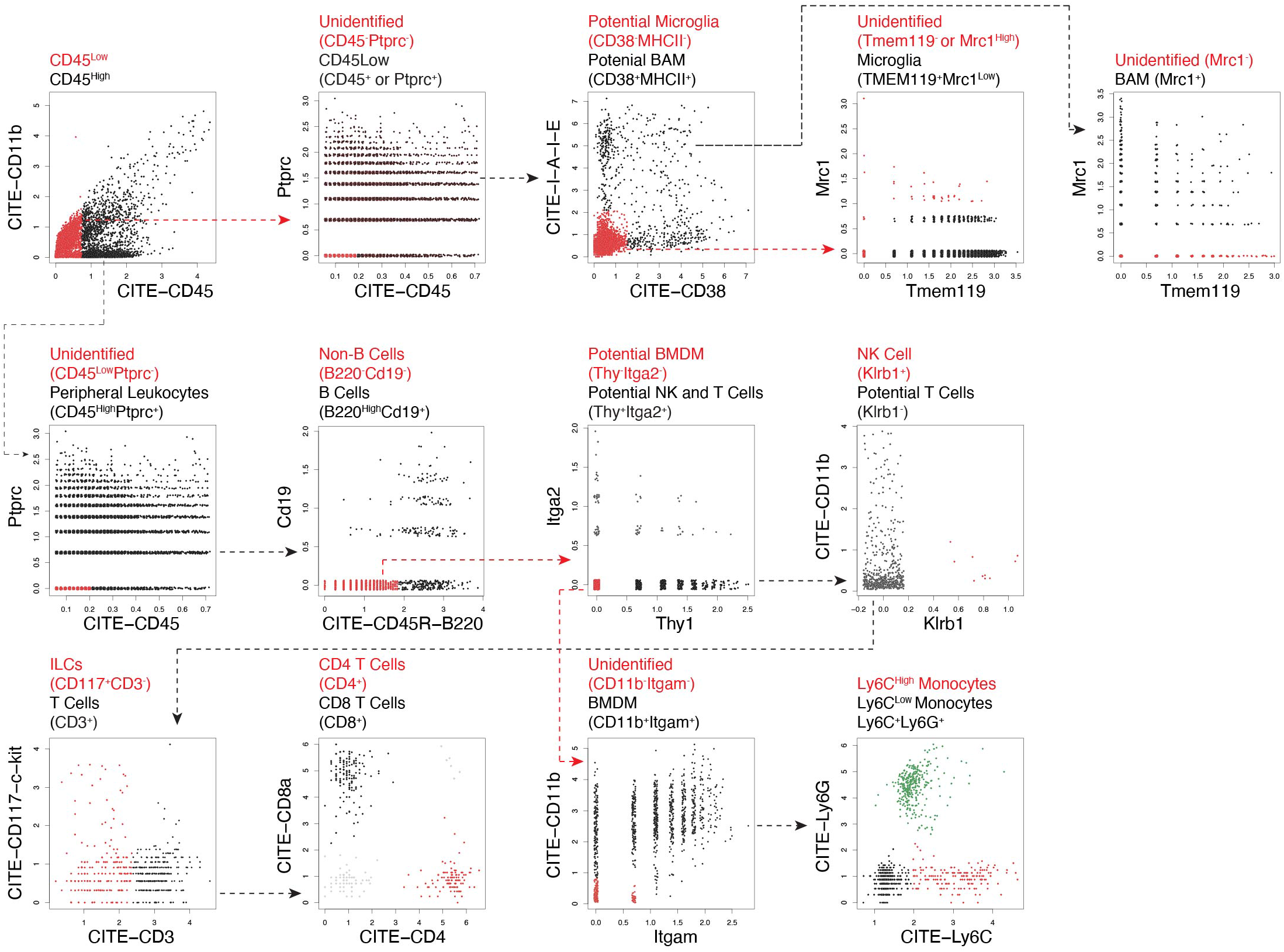
Gating strategy utilizing CITE antibodies and RNA to identify immune cell subsets in the brain.

**Supplementary Figure 3.**
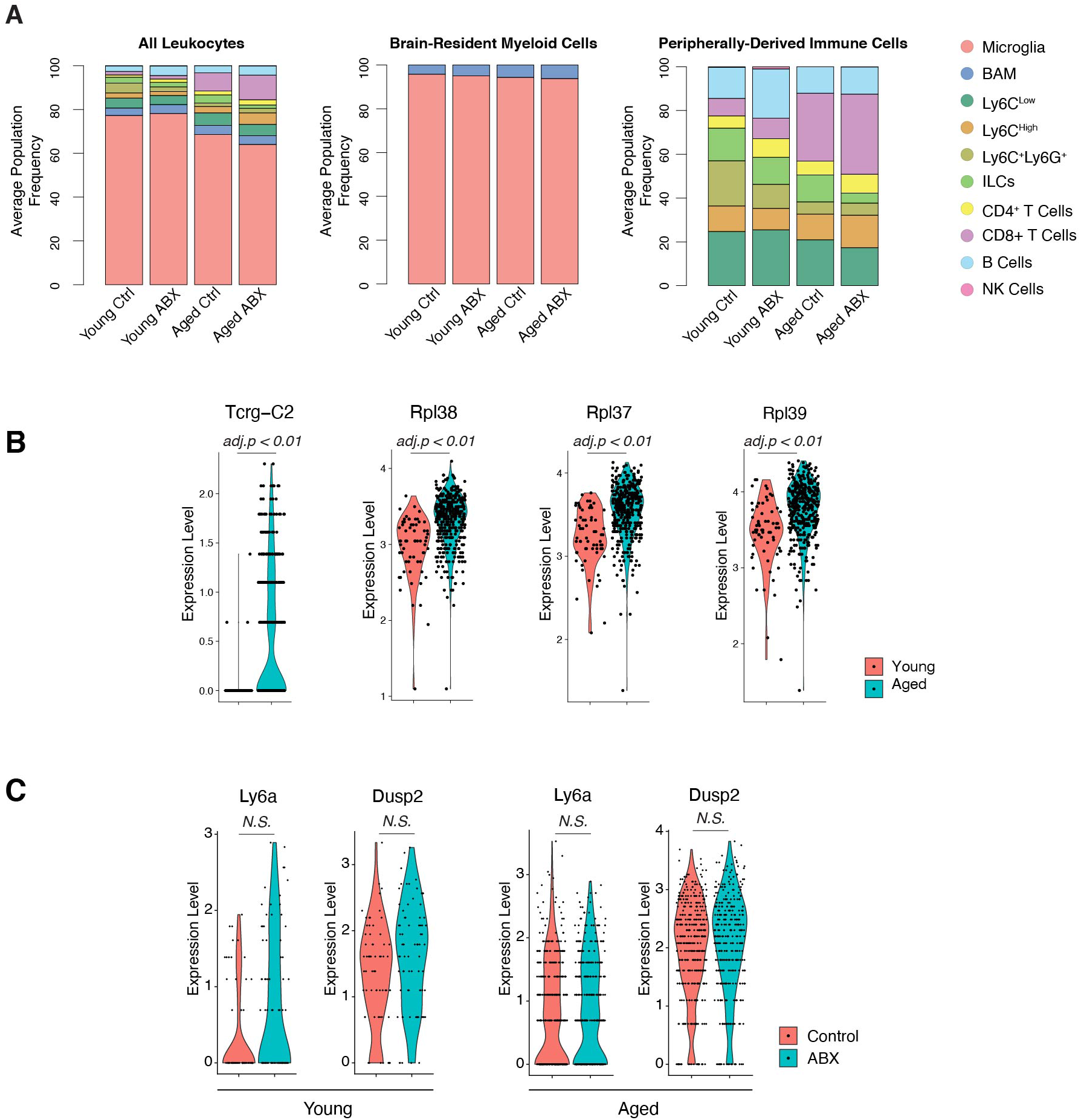
Cell-ID Proportions and Differentially Expressed Gene in CD8+ T cells. A) Stacked bar charts showing Cell-ID immune type frequencies compared against all leukocytes (left), brain-resident myeloid (middle) and peripherally-derived (right) immune populations. B) Violin plots showing upregulation of *Tcrg-C2* and ribosomal protein genes in CD8+ T cells from aged mice. C) Violin plots showing non-significant differential expression of Ly6a and *Dusp2* in CD8+ T cells from young (left) or aged (right) mice compared to their ABX treated counterparts.

**Supplementary Figure 4.**
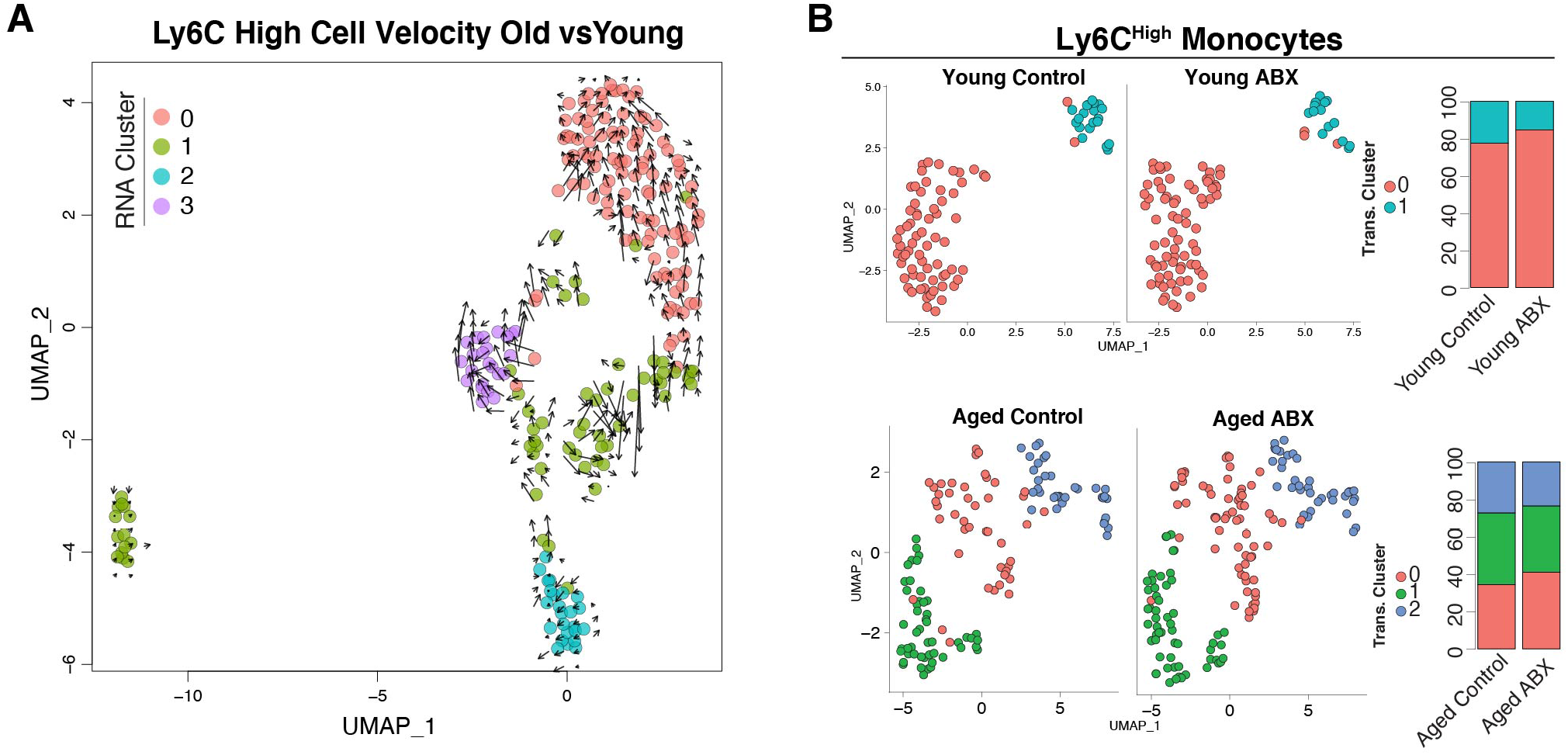
Ly6C^Hi^ cells response to ABX treatment. A) RNA velocity analysis showing the dynamics and interrelationship among subclusters Ly6C high monocytes. B) UMAP dimension plot showing the distribution of subcluster CNS Ly6C high cells in young or old brain with/without ABX treatment.

**Supplementary Figure 5.**
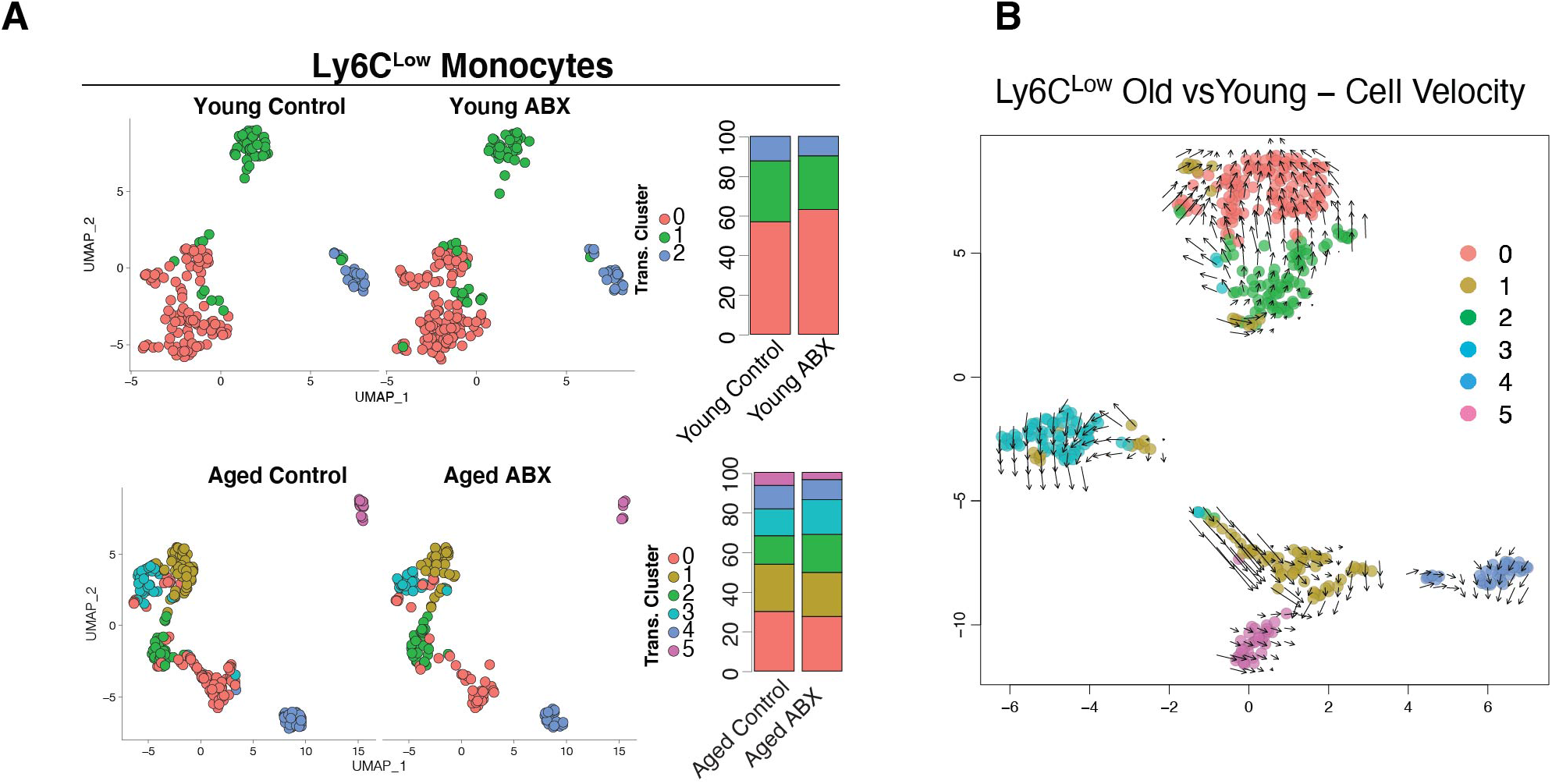
Ly6C^low^ cells response to ABX treatment. A) UMAP dimension plot showing the distribution of subcluster CNS Ly6C low cells in young or aged brain with/without ABX treatment. B) RNA velocity analysis showing the dynamics and interrelationship among subclusters Ly6C low monocytes.

**Supplementary Figure 6.**
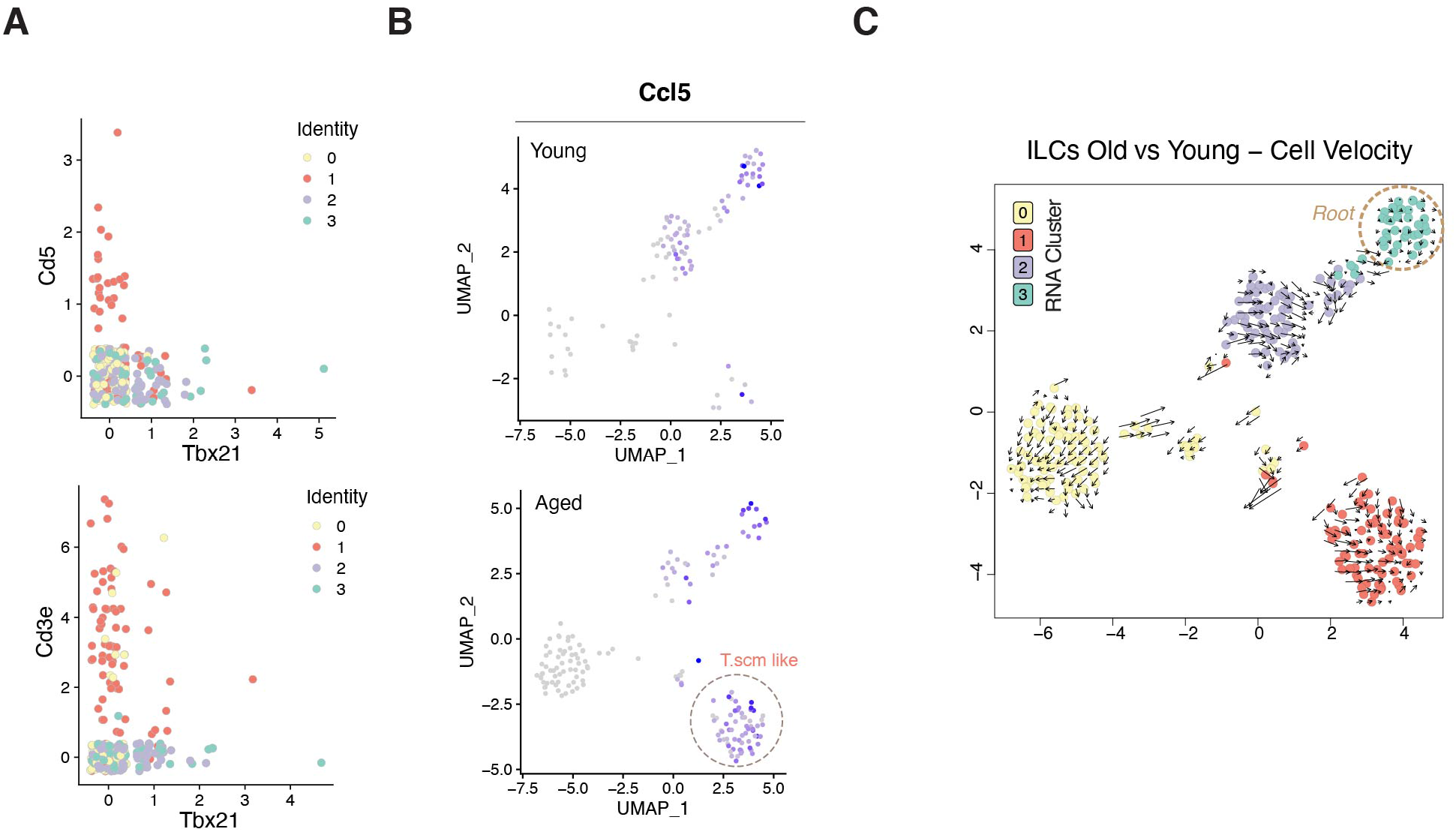
Gene expression of CNS ILCs and ILC RNA velocity analysis. A) Scatter plots showing the expression of T cell markers with Tbx21. B) UMAP dimention plot showing the distribution of Ccl5+ cells in the young or old brain. C) RNA velocity analysis showing the dynamics and interrelationship among subclusters CNS ILCs.

**Supplementary Figure 7.**
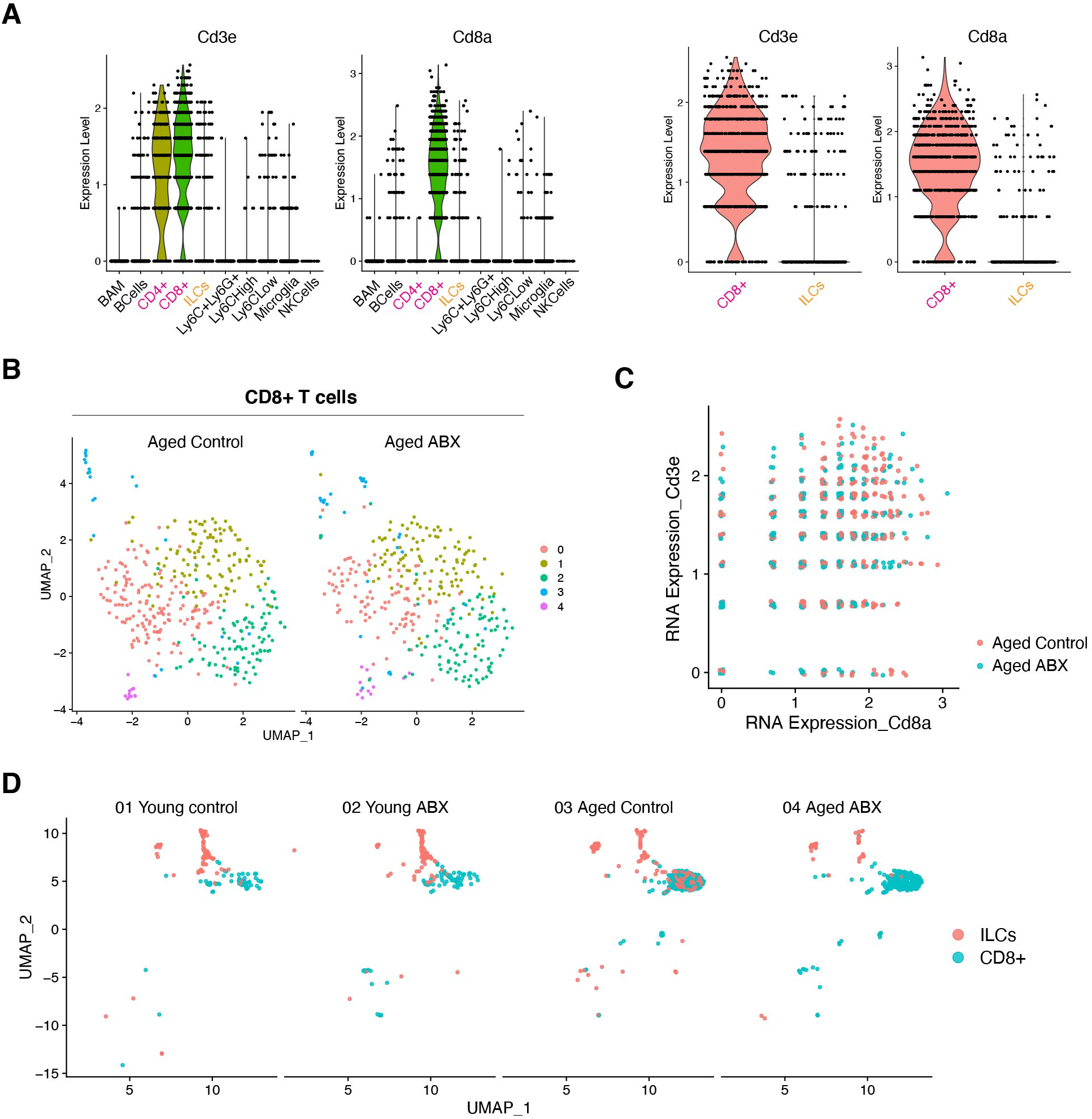
Comparison of classic CD8+ T cells and ILCs. A) Violin plots showing the relative RNA expression of Cd3e and Cd8a in Cell-ID ILCs and CD8+ T cells. B) UMAP feature plot showing overall changes of transcriptome of CD8+ T cells after ABX treatment in aged brain. C) Scatter plots showing the expression of T cell markers after ABX treatment in aged brain. D) UMAP dimension plot showing the distribution of Cell-ID ILCs and CD8+ T cells in the young or aged brains with/without ABX treatment.

## Notes

### Competing Interest Statement

The authors have declared no competing interest.

## REFERENCES

1. Al-Mossawi, H., Yager, N., Taylor, C.A., Lau, E., Danielli, S., de Wit, J., Gilchrist, J., Nassiri, I., Mahe, E.A., Lee, W., et al. (2019). Context-specific regulation of surface and soluble IL7R expression by an autoimmune risk allele. Nat. Commun. 10, 4575.

2. Arpaia, N., Campbell, C., Fan, X., Dikiy, S., van der Veeken, J., deRoos, P., Liu, H., Cross, J.R., Pfeffer, K., Coffer, P.J., et al. (2013). Metabolites produced by commensal bacteria promote peripheral regulatory T-cell generation. Nature 504, 451–455.

3. Auffray, C., Sieweke, M.H., and Geissmann, F. (2009). Blood monocytes: development, heterogeneity, and relationship with dendritic cells. Annu. Rev. Immunol. 27, 669–692.

4. Bachem, A., Makhlouf, C., Binger, K.J., de Souza, D.P., Tull, D., Hochheiser, K., Whitney, P.G., Fernandez-Ruiz, D., Dähling, S., Kastenmüller, W., et al. (2019). Microbiota-Derived Short-Chain Fatty Acids Promote the Memory Potential of Antigen-Activated CD8+ T Cells. Immunity 51, 285–297.e5.

5. Bal, S.M., Golebski, K., and Spits, H. (2020). Plasticity of innate lymphoid cell subsets. Nat. Rev. Immunol. 1–14.

6. Belkaid, Y., and Hand, T. (2014). Role of the Microbiota in Immunity and inflammation. Cell 157, 121–141.

7. Callahan, B.J., McMurdie, P.J., Rosen, M.J., Han, A.W., Johnson, A.J.A., and Holmes, S.P. (2016). DADA2: High-resolution sample inference from Illumina amplicon data. Nat. Methods 13, 581–583.

8. Colonna, M. (2018). Innate Lymphoid Cells: Diversity, Plasticity, and Unique Functions in Immunity. Immunity 48, 1104–1117.

9. Consortium, T.T.M., Pisco, A.O., Schaum, N., McGeever, A., Karkanias, J., Neff, N.F., Darmanis, S., Wyss-Coray, T., and Quake, S.R. (2019). A Single Cell Transcriptomic Atlas Characterizes Aging Tissues in the Mouse. BioRxiv 661728.

10. Derecki, N.C., Aleman-Muench, G.R., Lewis, G., Banie, H., Eckert, W., He, Y., Fourgeaud, L., Rao, S., Ma, J.Y., Carreira, V., et al. (2019). Meningeal Type-2 Innate Lymphoid Cells Emerge as Novel Regulators of Microglial Activation and Blood-Brain Barrier Stability: A Central Role for IL-10 (Rochester, NY: Social Science Research Network).

11. Dinan, T.G., and Cryan, J.F. (2017). Gut instincts: microbiota as a key regulator of brain development, ageing and neurodegeneration. J. Physiol. 595, 489–503.

12. Dulken, B.W., Buckley, M.T., Navarro Negredo, P., Saligrama, N., Cayrol, R., Leeman, D.S., George, B.M., Boutet, S.C., Hebestreit, K., Pluvinage, J.V., et al. (2019). Single-cell analysis reveals T cell infiltration in old neurogenic niches. Nature.

13. Ealey, K.N., and Koyasu, S. (2017). How Many Subsets of Innate Lymphoid Cells Do We Need? Immunity 46, 10–13.

14. Erickson, M.A., and Banks, W.A. (2019). Age-Associated Changes in the Immune System and Blood–Brain Barrier Functions. Int. J. Mol. Sci. 20.

15. Erny, D., Hrabĕ de Angelis, A.L., Jaitin, D., Wieghofer, P., Staszewski, O., David, E., Keren-Shaul, H., Mahlakoiv, T., Jakobshagen, K., Buch, T., et al. (2015). Host microbiota *constantly control maturation and function of microglia in the CNS*. Nat. Neurosci. 18, 965–977.

16. Franceschi, C., Garagnani, P., Parini, P., Giuliani, C., and Santoro, A. (2018). Inflammaging: a new immune-metabolic viewpoint for age-related diseases. Nat. Rev. Endocrinol. 14, 576–590.

17. Gagliani, N., Amezcua Vesely, M.C., Iseppon, A., Brockmann, L., Xu, H., Palm, N.W., de Zoete, M.R., Licona-Limón, P., Paiva, R.S., Ching, T., et al. (2015). Th17 cells transdifferentiate into regulatory T cells during resolution of inflammation. Nature 523, 221–225.

18. Galli, S.J., Borregaard, N., and Wynn, T.A. (2011). Phenotypic and functional plasticity of cells of innate immunity: macrophages, mast cells and neutrophils. Nat. Immunol. 12, 1035–1044.

19. Gattinoni, L., Speiser, D.E., Lichterfeld, M., and Bonini, C. (2017). T memory stem cells in health and disease. Nat. Med. 23, 18–27.

20. Gellhaar, S., Sunnemark, D., Eriksson, H., Olson, L., and Galter, D. (2017). Myeloperoxidase-immunoreactive cells are significantly increased in brain areas affected by neurodegeneration in Parkinson’s and Alzheimer’s disease. Cell Tissue Res. 369, 445–454.

21. Giri, M., Shah, A., Upreti, B., and Rai, J.C. (2017). Unraveling the genes implicated in Alzheimer’s disease. Biomed. Rep. 7, 105–114.

22. Gury-BenAri, M., Thaiss, C.A., Serafini, N., Winter, D.R., Giladi, A., Lara-Astiaso, D., Levy, M., Salame, T.M., Weiner, A., David, E., et al. (2016). The Spectrum and Regulatory Landscape of Intestinal Innate Lymphoid Cells Are Shaped by the Microbiome. Cell 166, 1231–1246.e13.

23. Jordão, M.J.C., Sankowski, R., Brendecke, S.M., Sagar, Locatelli, G., Tai, Y.-H., Tay, T.L., Schramm, E., Armbruster, S., Hagemeyer, N., et al. (2019). Single-cell profiling identifies myeloid cell subsets with distinct fates during neuroinflammation. Science 363, eaat7554.

24. Keren-Shaul, H., Spinrad, A., Weiner, A., Matcovitch-Natan, O., Dvir-Szternfeld, R., Ulland, T.K., David, E., Baruch, K., Lara-Astaiso, D., Toth, B., et al. (2017). A Unique Microglia Type Associated with Restricting Development of Alzheimer’s Disease. Cell 169, 1276–1290.e17.

25. Knox, J.J., Cosma, G.L., Betts, M.R., and McLane, L.M. (2014). Characterization of T-Bet and Eomes in Peripheral Human Immune Cells. Front. Immunol. 5.

26. Kwong, B., Rua, R., Gao, Y., Flickinger, J., Wang, Y., Kruhlak, M.J., Zhu, J., Vivier, E., McGavern, D.B., and Lazarevic, V. (2017). T-bet-dependent NKp46 + innate lymphoid *cells regulate the onset of T H 17-induced neuroinflammation*. Nat. Immunol. 18, 1117–1127.

27. Lang, R., and Raffi, F.A.M. (2019). Dual-Specificity Phosphatases in Immunity and Infection: An Update. Int. J. Mol. Sci. 20.

28. Langille, M.G., Meehan, C.J., Koenig, J.E., Dhanani, A.S., Rose, R.A., Howlett, S.E., and Beiko, R.G. (2014). Microbial shifts in the aging mouse gut. Microbiome 2, 50.

29. Lehallier, B., Gate, D., Schaum, N., Nanasi, T., Lee, S.E., Yousef, H., Moran Losada, P., Berdnik, D., Keller, A., Verghese, J., et al. (2019). Undulating changes in human plasma proteome profiles across the lifespan. Nat. Med. 25, 1843–1850.

30. Levy, M., Kolodziejczyk, A.A., Thaiss, C.A., and Elinav, E. (2017). Dysbiosis and the immune system. Nat. Rev. Immunol. 17, 219–232.

31. Ma, Q., Xing, C., Long, W., Wang, H.Y., Liu, Q., and Wang, R.-F. (2019). Impact of microbiota on central nervous system and neurological diseases: the gut-brain axis. J. Neuroinflammation 16, 53.

32. Manno, G.L., Soldatov, R., Zeisel, A., Braun, E., Hochgerner, H., Petukhov, V., Lidschreiber, K., Kastriti, M.E., Lönnerberg, P., Furlan, A., et al. (2018). RNA velocity of single cells. Nature 560, 494–498.

33. Meister, M., and Ferrandon, D. (2011). Immune cell transdifferentiation: a complex crosstalk between circulating immune cells and the haematopoietic niche. EMBO Rep. 13, 3–4.

34. Michaud, J.-P., Bellavance, M.-A., Préfontaine, P., and Rivest, S. (2013). Real-Time In Vivo Imaging Reveals the Ability of Monocytes to Clear Vascular Amyloid Beta. Cell Rep. 5, 646–653.

35. Mildner, A., Schönheit, J., Giladi, A., David, E., Lara-Astiaso, D., Lorenzo-Vivas, E., Paul, F., Chappell-Maor, L., Priller, J., Leutz, A., et al. (2017). Genomic Characterization of Murine Monocytes Reveals C/EBPβ Transcription Factor Dependence of Ly6C-Cells. Immunity 46, 849–862.e7.

36. Mrdjen, D., Pavlovic, A., Hartmann, F.J., Schreiner, B., Utz, S.G., Leung, B.P., Lelios, I., Heppner, F.L., Kipnis, J., Merkler, D., et al. (2018). High-Dimensional Single-Cell Mapping of Central Nervous System Immune Cells Reveals Distinct Myeloid Subsets in Health, Aging, and Disease. Immunity 48, 380–395.e6.

37. O’Toole, P.W., and Jeffery, I.B. (2015). Gut microbiota and aging. Science 350, 1214–1215.

38. Papalexi, E., and Satija, R. (2018). Single-cell RNA sequencing to explore immune cell heterogeneity. Nat. Rev. Immunol. 18, 35–45.

39. Plaza-Zabala, A., Sierra-Torre, V., and Sierra, A. (2017). Autophagy and Microglia: Novel Partners in Neurodegeneration and Aging. Int. J. Mol. Sci. 18.

40. Quail, D.F., and Joyce, J.A. (2017). The Microenvironmental Landscape of Brain Tumors. Cancer Cell 31, 326–341.

41. Romero-Suárez, S., Del Rio Serrato, A., Bueno, R.J., Brunotte-Strecker, D., Stehle, C., Figueiredo, C.A., Hertwig, L., Dunay, I.R., Romagnani, C., and Infante-Duarte, C. (2019). The Central Nervous System Contains ILC1s That Differ From NK Cells in the Response to Inflammation. Front. Immunol. 10, 2337.

42. Saelens, W., Cannoodt, R., Todorov, H., and Saeys, Y. (2019). A comparison of single-cell trajectory inference methods. Nat. Biotechnol. 37, 547–554.

43. Sampson, T.R., Debelius, J.W., Thron, T., Janssen, S., Shastri, G.G., Ilhan, Z.E., Challis, C., Schretter, C.E., Rocha, S., Gradinaru, V., et al. (2016). Gut Microbiota Regulate Motor Deficits and Neuroinflammation in a Model of Parkinson’s Disease. Cell 167, 1469–1480.e12.

44. Saresella, M., Marventano, I., Calabrese, E., Piancone, F., Rainone, V., Gatti, A., Alberoni, M., Nemni, R., and Clerici, M. (2014). A Complex Proinflammatory Role for Peripheral Monocytes in Alzheimer’s Disease. J. Alzheimers Dis. 38, 403–413.

45. Shen, X.Z., Okwan-Duodu, D., Blackwell, W.-L., Ong, F.S., Janjulia, T., Bernstein, E.A., Fuchs, S., Alkan, S., and Bernstein, K.E. (2014). Myeloid expression of angiotensin-converting enzyme facilitates myeloid maturation and inhibits the development of myeloid-derived suppressor cells. Lab. Investig. J. Tech. Methods Pathol. 94, 536–544.

46. Simoni, Y., and Newell, E.W. (2017). Toward Meaningful Definitions of Innate-Lymphoid-Cell Subsets. Immunity 46, 760–761.

47. Steinbach, K., Vincenti, I., Egervari, K., Kreutzfeldt, M., van der Meer, F., Page, N., Klimek, B., Rossitto-Borlat, I., Di Liberto, G., Muschaweckh, A., et al. (2019). Brain-resident memory T cells generated early in life predispose to autoimmune disease in mice. Sci. Transl. Med. 11.

48. Stoeckius, M., Hafemeister, C., Stephenson, W., Houck-Loomis, B., Chattopadhyay, P.K., Swerdlow, H., Satija, R., and Smibert, P. (2017). Simultaneous epitope and transcriptome measurement in single cells. Nat. Methods 14, 865–868.

49. Thaiss, C.A., Zmora, N., Levy, M., and Elinav, E. (2016). The microbiome and innate immunity. Nature 535, 65–74.

50. Thériault, P., ElAli, A., and Rivest, S. (2015). The dynamics of monocytes and microglia in Alzheimer’s disease. Alzheimers Res. Ther. 7, 41.

51. Ximerakis, M., Lipnick, S.L., Innes, B.T., Simmons, S.K., Adiconis, X., Dionne, D., Mayweather, B.A., Nguyen, L., Niziolek, Z., Ozek, C., et al. (2019). Single-cell transcriptomic profiling of the aging mouse brain. Nat. Neurosci. 22, 1696–1708.

52. Yang, J., Zhang, L., Yu, C., Yang, X.-F., and Wang, H. (2014). Monocyte and macrophage differentiation: circulation inflammatory monocyte as biomarker for inflammatory diseases. Biomark. Res. 2, 1.

53. Zaiss, D.M.W., Gause, W.C., Osborne, L.C., and Artis, D. (2015). Emerging Functions of Amphiregulin in Orchestrating Immunity, Inflammation, and Tissue Repair. Immunity 42, 216–226.

54. Zhou, X., Liao, W.-J., Liao, J.-M., Liao, P., and Lu, H. (2015). Ribosomal proteins: functions beyond the ribosome. J. Mol. Cell Biol. 7, 92–104.

